# Reconstruction Algorithms for DNA-Storage Systems

**DOI:** 10.1101/2020.09.16.300186

**Authors:** Omer Sabary, Alexander Yucovich, Guy Shapira, Eitan Yaakobi

**Affiliations:** Computer Science Department, Technion, Haifa, 3200003, Israel

## Abstract

In the *trace reconstruction problem* a length-*n* string ***x*** yields a collection of noisy copies, called *traces*, ***y***_1_, …, ***y***_*t*_ where each ***y***_*i*_ is independently obtained from ***x*** by passing through a *deletion channel*, which deletes every symbol with some fixed probability. The main goal under this paradigm is to determine the required minimum number of i.i.d traces in order to reconstruct ***x*** with high probability. The trace reconstruction problem can be extended to the model where each trace is a result of ***x*** passing through a *deletion-insertion-substitution channel*, which introduces also insertions and substitutions. Motivated by the storage channel of DNA, this work is focused on another variation of the trace reconstruction problem, which is referred by the *DNA reconstruction problem*. A *DNA reconstruction algorithm* is a mapping 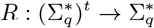 which receives *t* traces ***y***_1_, …, ***y***_*t*_ as an input and produces 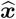, an estimation of ***x***. The goal in the DNA reconstruction problem is to minimize the edit distance 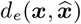 between the original string and the algorithm’s estimation. For the deletion channel case, the problem is referred by the *deletion DNA reconstruction problem* and the goal is to minimize the Levenshtein distance 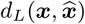.

In this work, we present several new algorithms for these reconstruction problems. Our algorithms look globally on the entire sequence of the traces and use dynamic programming algorithms, which are used for the *shortest common supersequence* and the *longest common subsequence* problems, in order to decode the original sequence. Our algorithms do not require any limitations on the input and the number of traces, and more than that, they perform well even for error probabilities as high as 0.27. The algorithms have been tested on simulated data as well as on data from previous DNA experiments and are shown to outperform all previous algorithms.

## 1 Introduction

Recent studies presented a significant progress in DNA synthesis and sequencing technologies [3, 11, 31, 32, 33, 41, 48]. This progress also introduced the development of data storage technology based upon DNA molecules. A DNA storage system consists of three important components. The first is the *DNA synthesis* which produces the *oligonucleotides*, also called *strands*, that encode the data. In order to produce strands with acceptable error rates, in a high throughput manner, the length of the strands is typically limited to no more than 250 nucleotides [5]. The second part is a storage container with compartments which stores the DNA strands, however without order. Finally, *sequencing* is performed to read back a representation of the strands, which are called *reads*.

Current synthesis technologies are not able to generate a single copy for each DNA strand, but only multiple copies where the number of copies is in the order of thousands to millions. Moreover, sequencing of DNA strands is usually preceded by PCR amplification which replicates the strands [25]. Hence, every strand has multiple copies and several of them are read during sequencing.

The encoding and decoding stages are two processes, external to the storage system, that convert the user’s binary data into strands of DNA such that, even in the presence of errors, it will be possible to revert back and recover the original binary data. These two stages consist of three steps, which we refer by 1. *clustering*, 2. *reconstruction*, and finally 3. *error correction*. After the strands are read back by sequencing, the first task is to partition them into *clusters* such that all strands in the same cluster originated from the same synthesized strand. After the clustering step, the goal is to reconstruct each strand based upon all its noisy copies, and this stage is the main problem studied in this paper. Lastly, errors which were not corrected by the reconstruction step, mis-clustering errors, lost strands, and any other error mechanisms should be corrected by the use of an error-correcting code.

Any reconstruction algorithm for the second stage is performed on each cluster to recover the original strand from the noisy copies in the cluster. Having several copies for each strand is beneficial since it allows to correct errors that may occur during this process. In fact, this setup falls under the general framework of the *string reconstruction problem* which refers to recovering a string based upon several noisy copies of it. Examples for this problem are the *sequence reconstruction problem* which was first studied by Levenshtein [34, 35] and the *trace reconstruction problem* [4, 16, 26, 27, 43]. In general, these models assume that the information is transmitted over multiple channels, and the decoder, which observes all channel estimations, uses this inherited redundancy in order to correct the errors.

Generally speaking, the main problem studied under the paradigm of the sequence reconstruction and trace reconstruction problems is to find the minimum number of channels that guarantee successful decoding either in the worst case or with high probability. However, in DNA-based storage systems we do not necessary have control on the number of strands in each cluster. Hence, the goal of this work is to propose efficient algorithms for the reconstruction problem as it is reflected in DNA-based storage systems where the cluster size is a given parameter. Then, the goal is to output a strand that is close to the original one so that the number of errors the error-correcting code should correct will be minimized. We will present algorithms that work with a flexible number of copies and various probabilities for deletion, insertion, and substitution errors.

In our model we assume that the clustering step has been done successfully. This could be achieved by the use of indices in the strands and other advanced coding techniques; for more details see [47] and references therein. Thus, the input to the algorithms is a cluster of noisy read strands, and the goal is to efficiently output the original strand or a close estimation to it with high probability. We also apply our algorithms on data from previously published DNA-storage experiments [20, 24, 40] and compare our accuracy and performance with state of the art algorithms known from the literature.

### 1.1 DNA Storage and DNA Errors

One of the early experiments of data storage in DNA was conducted by Clellan et al. in 1999. In their study they coded and recovered a message consisting of 23 characters [15]. Three sequences of nine bits each, have been successfully stored by Leier et al. in 2000. Gibson et al. [21] presented in 2010 a more significant progress, in terms of the amount of data stored successfully. They demonstrated in-vivo storage of 1,280 characters in a bacterial genome. The first large scale demonstrations of the potential of in vitro DNA storage were reported by Church et al. who recovered 643 KB of data [14] and by Goldman et al. who accomplished the same task for a 739 KB message [22]. However, both of these pioneering groups did not recover the entire message successfully and no error correcting codes were used. Shortly later, in [24], Grass et al. have managed to successfully store and recover a 81 KB message, in an encapsulated media, and Bornholt et al. demonstrated storing a 42 KB message [7]. A significant improvement in volume was reported in [6] by Blawat et al. who successfully stored 22 MB of data. Erlich and Zielinski improved the storage density and stored 2.11 MB of data [20]. The largest volume of stored data was reported by Organick et al. in [40] who stored roughly 200 MB of data, an order of magnitude more data than previously reported. Yazdi et al. developed in [54] a method that offers both random access and rewritable storage and in [53] a portable DNA-based storage system. Recently, Anavy et al. [1] enhanced the capacity of the DNA storage channel by using composite DNA letter. A similar approach, on a smaller scale, was reported in [13]. Lopez et al. stored and decoded a 1.67 MB of data in [36]. In their work they focused on increasing the throughput of nanopore sequencing by assembling and sequencing together fragments of 24 short DNA strands. Recent studies also presented an end-to-end demonstration of DNA storage [51], the use of LDPC codes for DNA-based storage [10], a computer systems prospective on molecular processing and storage [9], and lastly, the work of Tabatabaei et al. [50] which uses existing DNA strands as punch cards to store information.

The processes of synthesizing, storing, sequencing, and handling strands are all error prone. Each step in these processes can independently introduce a significant number of errors. Additionally, the DNA storage channel has several attributes which distinguish it from other storage media such as tapes, hard disk drives, and flash memories. We summarize some of these differences and the special error behavior in DNA.

1. Both the synthesis and sequencing processes can introduce deletion, insertion, and substitution errors on each of the read and synthesized strands.
2. Current synthesis methods cannot generate one copy for each design strand. They all generate thousands to millions of noisy copies, while different copies may have a different error distribution. Moreover, some strands may have a significant larger number of copies, while some other strands may not have copies all.
3. The use of DNA for storage or other applications typically involves PCR amplification of the strands in the DNA pool [25]. PCR is known to have a preference for some strands over others, which may further distort the distribution of the number of copies of individual strands and their error profiles [42, 44].
4. Longer DNA strands can be sequenced using designated sequencing technologies, e.g. PacBio and Oxford Nanopore [10, 36, 40, 53]. However, the error rates of these technologies can be significantly higher and can grow up to 30%, with deletions and substitutions as the most dominant errors [45].

A detailed characterization of the errors in the DNA-storage channel and more statistics about previous DNA-storage experiments can be found in [25, 45].

### 1.2 This Work

In this work we present several reconstruction algorithms, crafted for DNA storage systems. Since our purpose is to solve the reconstruction problem as it is reflected in DNA-storage systems, our algorithms aim to minimize the distance between the output and the original strands. The algorithms in this work are different from most of the previously published reconstruction algorithms in several respects. Firstly, we do not require any assumption on the input. That is, the input can be arbitrary and does not necessarily belong to an error-correcting code. Secondly, our algorithms are not limited to specific cluster size, do not require any dependencies between the error probabilities, and do not assume zero errors in any specific location of the strands. Thirdly, we limit the complexity of our algorithms, so they can run with practical time on actual data from previous DNA-storage experiments. Lastly, since clusters in DNA storage systems may vary in their size and errors distributions, our algorithms are designed to minimize the distance between our output and the original strand, taking into account these errors can be corrected by the use of an error-correcting code.

## 2 Preliminaries and Problem Definition

We denote by Σ_*q*_ = {0, …, *q − 1*} the alphabet of size *q* and 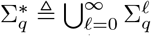. The length of ***x*** *∈* Σ^*n*^ is denoted by |***x***| = *n*. The *Levenshtein distance* between two strings 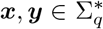, denoted by *d*_*L*_(***x, y***), is the minimum number of insertions and deletions required to transform ***x*** into ***y***. The *edit distance* between two strings 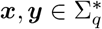, denoted by *d*_*e*_(***x, y***), is the minimum number of insertions, deletions and substitution required to transform ***x*** into ***y***, and *d*_*H*_ (***x, y***) denotes the *Hamming distance* between ***x*** and ***y***, when |***x***| = |***y***|. For a positive integer *n*, the set {1, …, *n*} is denoted by [*n*].

### 2.1 The DNA Reconstruction Problem

The *trace reconstruction problem* was first proposed in [4] and was later studied in several theoretical works; see e.g. [26, 27, 39, 43]. Under this framework, a length-*n* string ***x***, yields a collection of noisy copies, also called *traces*, ***y***_1_, ***y***_2_, …, ***y***_*t*_ where each ***y***_*i*_ is independently obtained from ***x*** by passing through a *deletion channel*, under which each symbol is independently deleted with some fixed probability *p*_*d*_. Suppose the input string ***x*** is arbitrary. In the trace reconstruction problem, the main goal is to determine the required minimum number of i.i.d traces in order to reconstruct ***x*** with high probability. This problem has two variants: in the “worst case”, the success probability refers to all possible strings, and in the “average case” (or “random case”) the success probability is guaranteed for an input string ***x*** which is chosen uniformly at random.

The trace reconstruction problem can be extended to the model where each trace is a result of ***x*** passing through a *deletion-insertion-substitution channel*. Here, in addition to deletions, each symbol can be switched with some substitution probability *p*_*s*_, and for each *j*, with probability *p*_*i*_, a symbol is inserted before the *j*-th symbol of ***x***^1^. Under this setup, the goal is again to find the minimum number of channels which guarantee successful reconstruction of ***x*** with high probability.

Motivated by the storage channel of DNA and in particular the fact that different clusters can be of different sizes, this work is focused on another variation of the trace reconstruction problem, which is referred by the *DNA reconstruction problem*. The setup is similar to the trace reconstruction problem. A length-*n* string ***x*** is transmitted *t* times over the *deletion-insertion-substitution channel* and generates *t* traces ***y***_1_, ***y***_2_, …, ***y***_*t*_. A *DNA reconstruction algorithm* is a mapping 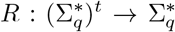 which receives the *t* traces ***y***_1_, ***y***_2_, …, ***y***_*t*_ as an input and produces 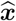, an estimation of ***x***. The goal in the DNA reconstruction problem is to minimize 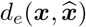, i.e., the edit distance between the original string and the algorithm’s estimation. When the channel of the problem is the *deletion channel*, the problem is referred by the *deletion DNA reconstruction problem* and the goal is to minimize 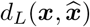. While the main figure of merit in these two problems is the edit/Levenshtein distance, we will also be concerned with the complexity, that is, the running time of the proposed algorithms.

## 3 Related Work

This section reviews the related works on the different reconstruction problems. In particular we list the reconstruction algorithms that have been used in previous DNA storage experiments and summarize some of the main theoretical results on the trace reconstruction problem.

### 3.1 Reconstruction Algorithms for DNA-Storage Systems

1. Baku et al. [4] studied the trace reconstruction problem as an abstraction and a simplification of the multiple sequence alignment problem in bioinformatics. Here the goal is to reconstruct the DNA of a common ancestor of several organisms using the genetic sequences of those organisms. They focused on the deletion case of this problem and suggested a majority-based algorithm to reconstruct the sequence, which they referred by the *bitwise majority alignment* (*BMA*) *algorithm*. They aligned all traces by considering the majority vote per symbol from all traces, while maintaining pointers for each of the traces. If a certain symbol from one (or more) of the traces does not agree with the majority symbol, its pointer is not incremented and it is considered as a deletion. They showed and proved that even though this technique works locally for each symbol, its success probability is relatively high when the deletion probability is small enough.
2. Viswanathan and Swaminathan presented in [52] a BMA-based algorithm for the trace reconstruction problem under the deletion-insertion-substitution channel. Their algorithm extends the BMA algorithm so it can support also insertions and substitutions. It works iteratively on “segments” from the traces, where a segment consists of consecutive bits and its size is a fixed fraction of the trace that is given as a parameter to the algorithm. The segment of each trace is defined by its pointers. The pointers of the traces are updated in each iteration similarly as in the BMA algorithm. Each trace is classified as valid or invalid by its distance from the majority segment. Once less than 3*/*4 of the traces’ segments are valid, the rest of the bits are estimated by the valid traces. Their algorithm extends [4] and improves the results from [29], where it was shown that when the number of traces is *O*(log *n*) and the deletion/insertion probability is 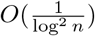, it is possible to reconstruct a sequence with high probability. However, it assumes the deletion and insertion probabilities are relatively small, while the substitution probability is relatively large. In practice, these probabilities vary from cluster to cluster and do not necessarily meet these assumptions.
3. Gopalan et al. [23] used the approach of the BMA algorithm from [4] and extended it to work with deletions, insertions, and substitutions to support DNA storage systems. They also considered a majority vote per symbol with some improvements. For any trace that its current symbol did not match the majority symbol, they used a “lookahead window” to look on the next 2 (or more) symbols. Then, they compared the next symbols to the majority symbols and classified it as an error accordingly. Organick et al. conducted a large scale DNA storage experiments in [40] where they successfully reconstructed their sequences using the reconstruction algorithm of Gopalan et al. [23].
4. For the case of sequencing via nanopore technology, Duda et al. [17] studied the trace reconstruction problem, while considering insertions, deletions, and substitutions. They focused on dividing the sequence into homopolymers (consecutive replicas of the same symbol), and proved that the number of copies required for accurately reconstructing a long strand is logarithmic with the strand’s length. Yazdi et al. used in [53] a similar but different approach in their DNA storage experiment. They first aligned all the strands in the cluster using the multiple sequence alignment algorithm MUSCLE [18, 37]. Then, they divided each strand into homopolymers and performed majority vote to determine the length of each homopolymer separately. Their strands were designed to be balanced in their GC-content, which means that 50% of the symbols in each strands were G or C. Hence, they could perform additional majority iterations on the homopolymers’ lengths until the majority sequence was balanced in its GC-content. All of these properties guaranteed successful reconstruction of the strands and therefore they did not need to use any error-correcting code in their experiment [53].

### 3.2 Theoreitical Results on the Trace Reconstruction Problem

1. Holenstein et al. [27] presented an algorithm for reconstructing a random string using polynomially many traces from the deletion channel. In their work they assumed that the deletion probability is constant and is smaller than some threshold *γ*. Their work suggested a slightly different technique than [4], in the sense that they did not use standard majority voting, but a different majority scheme, where each trace vote is utilized with a probability measure of its certainty. They assumed that the last *O*(log *n*) of the input string (“anchor”) can not be affected by the deletion channel, or be removed to another part of the strings. Thus, the certainty of each trace is estimated by checking if the last *O*(log *n*) bits of the trace (“anchor”) match the last *O*(log *n*) of the recovered string. The threshold *γ* from [27] was later estimated to be at most 0.07 by Peres and Zhai [43].
2. Peres and Zhai [43] also improved the work by Holenstein et al. [27] and the work by McGregor et al. [38]. They not only extended the range of the supported deletion probability to be [0, 1*/*2), but also showed that a subpolynomial number of traces, more specifically exp(*O*(log^1*/*2^ *n*)), is sufficient for the reconstruction of a random string. Their approach includes two steps, where in the first one the strings are aligned and then, in the second step, a majority-based algorithm is invoked in order to reconstruct the sequence. The alignment step of their algorithm is quite similar to the “anchor” technique as presented in [27]. The difference is that Peres and Zhai did not restrict the “anchor” to be just the last *O*(log *n*) bits in the strings, but it could be placed at any position in the traces.
3. Holden, Pemantle, and Peres improved in [26] the upper bound on the number of traces for the random case to (exp *O*(log^1*/*3^ *n*)) while both insertions and deletions are allowed in any probability from the range [0, 1). Their algorithm consists of three ingredients: (i) A boolean test *T* (*w, w*^*′*^) works on pairs of sequences of the same length and indicates whether *w*^*′*^ is likely to be a result of *x* passing through the deletion-insertion channel. (ii) An alignment procedure that creates for each of the traces *y*_*i*_ an estimate *τ* for the matching between positions in *y*_*i*_ and *x*. (iii) A bit recovery procedure that uses the aligned traces in order to estimate a bit or subsequence of bits.
4. For the worst case scenario, in [27] it is shown that exp *O*(*n*^1*/*2^ log *n*) traces suffice for reconstruction with high probability. This was later improved independently by both De, O’Donnell, and Severdio in [16] and by Nazarov and Peres [39] to exp *O*(*n*^1*/*3^).
5. Two recent works [8, 12] studied the trace reconstruction problem for the codes setup, i.e., the transmitted is sequence not arbitrary but belongs to some code with error-correction capabilities.
6. Another related model has been studied by Kiah et al. who introduced in [30] another approach for the trace reconstruction problem, where they used profile-vectors-based coding scheme in order to reconstruct the sequence.
7. Another related problem, phrased as the *sequence reconstruction problem*, was also studied by Levenshtein in [34] and [35], but his approach was different. Under this paradigm he studied the minimum number of different (noisy) channels that is required in order to build a decoder that can reconstruct any transmitted sequence in the worst case. In [34], he showed that the number of channels that is required to recover any sequence has to be greater than the maximum intersection between the error balls of any two transmitted sequences. Since Levenshtein studied the worst case of this problem, the number of unique channels has to be extremely large which is not applicable for the practical setup we consider in the reconstruction of DNA strands in a DNA-based storage system.

## 4 Supersequences, Subsequences, and Maximum Likelihood

While all previous works of reconstructing algorithms used variations of the majority algorithm on localized areas of the traces, we take a different global approach to tackle this problem. Namely, the algorithms presented in the paper are heavily based on the *maximum likelihood decoder* for multiple deletion channels as studied recently in [46, 49] as well as the concepts of the *shortest common supersequence* and the *longest common subsequence*. Hence, we first briefly review the main ideas of these concepts, while the reader is referred to [46] for a more comprehensive study of this summary.

### 4.1 Supersequences and Subsequences

For a sequence 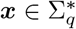 and a set of indices *I* ⊆ [|***x***|], the sequence ***x***_*I*_ is the *projection* of ***x*** on the indices of *I* which is the subsequence of ***x*** received by the symbols at the entries of *I*. A sequence ***x*** *∈* Σ^*\**^ is called a *supersequence* of ***y*** *∈* Σ^*\**^, if ***y*** can be obtained by deleting symbols from ***x***, that is, there exists a set of indices *I ⊆* [|***x***|] such that ***y*** = ***x***_*I*_. In this case, it is also said that ***y*** is a *subsequence* of ***x***. Furthermore, ***x*** is called a *common supersequence* (*subsequence*) of some sequences ***y***_1_, …, ***y***_*t*_ if ***x*** is a supersequence (subsequence) of each one of these *t* sequences. The *length of the shortest common supersequence* (*SCS)* of ***y***_1_, …, ***y***_*t*_ is denoted by SCS(***y***_1_, …, ***y***_*t*_). The set of all shortest common supersequences of ***y***_1_, …, ***y***_*t*_ *∈* Σ^*\**^ is denoted by *𝒮*CS(***y***_1_, …, ***y***_*t*_). Similarly, the *length of the longest common subsequence* (*LCS)* of ***y***_1_, …, ***y***_*t*_, is denoted by LCS(***y***_1_, …, ***y***_*t*_), and the set of all longest subsequences of ***y***_1_, …, ***y***_*t*_ is denoted by *ℒ*CS(***y***_1_, …, ***y***_*t*_).

### 4.2 Maximum Likelihood Decoder for Multiple Deletion Channels

Consider a channel S that is characterized by a conditional probability Pr_S_, which is defined by

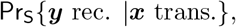

for every pair 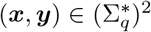. Note that it is not assumed that the lengths of the input and output sequences are the same as we consider also deletions and insertions of symbols. As an example, it is well known that if S is the *binary symmetric channel* (*BSC*) with crossover probability 0 ⩽ *p* ⩽ 1*/*2, denoted by BSC(*p*), it holds that 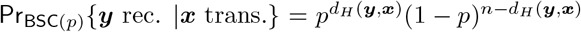, for all 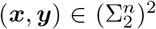, and otherwise (the lengths of ***x*** and ***y*** is not the same) this probability equals 0.

The *maximum-likelihood* (*ML*) *decoder* for a code *𝒞* with respect to S, denoted by *𝒟*_ML_, outputs a codeword ***c*** *∈ 𝒞* that maximizes the probability Pr_S_{****y*** rec. |***c*** trans.*}. That is, for 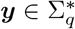,

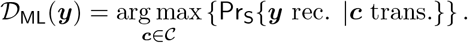

It is well known that for the BSC, the ML decoder simply chooses the closest codeword with respect to the Hamming distance.

The conventional setup of channel transmission is extended to the case of more than a single instance of the channel. Assume a sequence ***x*** is transmitted over some *t* identical channels of S and the decoder receives all channel outputs ***y***_1_, …, ***y***_*t*_. This setup is characterized by the conditional probability

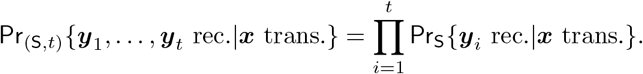

Now, the input to the ML decoder is the sequences ***y***_1_, …, ***y***_*t*_ and the output is the codeword ***c*** which maximizes the probability Pr_(S,*t*)_{****y***_1_, …, ***y***_*t*_ rec.|***x*** trans.*}.

For two sequences 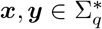, the number of times that ***y*** can be received as a subsequence of ***x*** is called the *embedding number of* ***y*** *in* ***x*** and is defined by

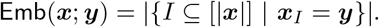

Note that if ***y*** is not a subsequence of ***x*** then Emb(***x***; ***y***) = 0. The embedding number has been studied in several previous works; see e.g. [2, 19] and in [49] it was referred to as the *binomial coefficient*. In particular, this value can be computed with quadratic complexity [19].

While the calculation of the conditional probability Pr_S_ {****y*** rec. |***x*** trans*}. is a rather simple task for many of the known channels, it is not straightforward for channels which introduce insertions and deletions. In the *deletion channel* with deletion probability *p*, denoted by Del(*p*), every symbol of the word ***x*** is deleted with probability *p*. For the deletion channel it is known, see e.g. [46, 49], that for all 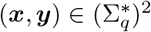, it holds that

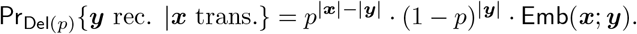

According to this property, the ML decoder for one or multiple deletion channels is stated as follows [46].

**Lemma 1**. *Assume* ***c*** *∈ 𝒞 ⊆* (Σ_*q*_)^*n*^ *is the transmitted sequence and* ***y***_1_, …, ***y***_*t*_ (Σ_*q*_)^*\**^ *are the output sequences from* Del(*p*), *then*

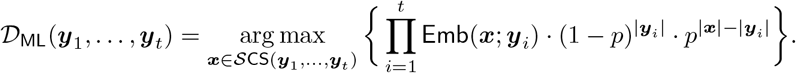

Note that since there is more than a single channel, when the goal is to minimize the average decoding error probability, the ML decoder does not necessarily have to output a codeword but any sequence that minimizes the average decoding error probability. In the next sections it will be shown how to use the concepts of the SCS and LCS together with the maximum likelihood decoder in order to build decoding algorithms for the deletion DNA reconstruction and the DNA reconstruction problems.

## 5 The Deletion DNA Reconstruction Problem

This section studies the deletion DNA reconstruction problem. Assume that a cluster consists of *t* traces, ***y***_1_, ***y***_2_, …, ***y***_*t*_, where all of them are noisy copies of a synthesized strand. This model assumes that every strand is a sequence that is independently received by the transmission of a length-*n* sequence ***x*** (the synthesized strand) through a deletion channel with some fixed deletion probability *p*_*d*_. Our goal is to propose an efficient algorithm which returns 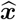, an estimation of the transmitted sequence ***x***, with the intention of minimizing the Levenshtein distance between ***x*** and 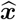. We consider both cases when *t* is a fixed small number and large values of *t*.

### 5.1 An Algorithm for Small Fixed Values of *t*

Our approach is based on the maximum likelihood decoder over the deletion channel as presented in [46, 49]. A straightforward implementation of this approach on a cluster of size *t* is to compute the set of shortest common supersequences of ***y***_1_, ***y***_2_, …, ***y***_*t*_, i.e., the set *S*CS(***y***_1_, ***y***_2_, …, ***y***_*t*_), and then return the maximum likelihood sequence among them. This algorithm has been rigorously studied in [46] to analyze its Levenshtein error rate for *t* = 2. The method to calculate the length of the SCS commonly uses dynamic programming [28] and its complexity is the product of the lengths of all sequences. Hence, even for moderate cluster sizes, e.g. *t* ⩾ 5, this solution will incur high complexity and impractical running times. However, for many practical values of *n* and *p*_*d*_, the original sequence ***x*** can be found among the list of SCSs while taking less than *t* traces or even only two of them. This fact, which we verified empirically, can drastically reduce the complexity of the ML-based algorithm. Furthermore, note that ***x*** is always a common supersequence of all traces, however it is not necessarily the shortest one. Hence, our algorithm works as follows. The algorithm creates sorted sets of *r*-tuples, where each tuple consists of *r* traces from the cluster. The *r*-tuples are sorted in a non-decreasing order according to the sum of their lengths. For each *r*-tuple 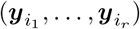, the algorithm first calculates its length of the SCS, i.e., the value 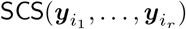. Observe that if 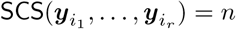 then the sequence ***x*** necessarily appears in the set of SCSs of 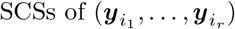, that is, *x* ∈ 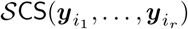. However it is not necessarily the only sequence in 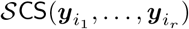. Hence, all is left to do is to filter the set 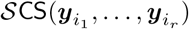 with sequences that are supersequences of all *t* traces and finally return the maximum likelihood among them. The algorithm iterates over all possible *r*-tuples for *r* = 2, 3, 4 and if none of them succeeds, the algorithm computes all SCSs of maximal length that were observed throughout its run and returns the one that minimizes the sum of Levenshtein distances from all copies in the cluster.

In Algorithm 1, we present a pseudo-code of our solution for the deletion DNA reconstruction problem. Note that the algorithm uses another procedure which is presented in Algorithm 2 to filter the supersequences and output the maximum likelihood supersequence. The input to the algorithm is the length *n* of the original sequence, and a cluster of *t* traces **C**. Algorithm 1’s main loop is in Step 2; first in Step 2-a it generates the set *F*, which is a sorted set of all *r*-tuples of traces by the sum of their lengths. Then, in Step 2-b it iterates over all *r*-tuples in *F* and checks for each *r*-tuple, 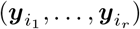, if the length of their SCS, i.e., 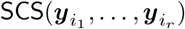, equals *n*. If it is equal to *n*, it computes the set of all its SCSs, 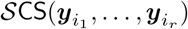, and invokes Algorithm 2. Algorithm 2 checks if one or more of those SCSs are supersequences of all of the traces in the cluster, and if so it returns the maximum likelihood among them. In case that 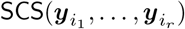, the algorithm checks also if it is equal or greater than *n*_max_, which is the longest SCS that was found so far. In this case, the algorithm saves **C**_max_, which is the set of all *r*-tuples such that the length of their SCS equals *n*_max_. In Step 3, the algorithm computes 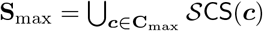, which is the union of sets of SCSs of the *r*-tuples that the length of their SCS was *n*_max_. In Step 4, the algorithm invokes again Algorithm 2 to check if **S**_max_ includes supersequences of all traces in **C** and returns the maximum likelihood among them. If none of the sequences in **S**_max_ is a supersequence of all traces in **C**, the algorithm returns in Step 5 the sequence which minimizes the sum of Levenshtein distances to all the traces in **C**.

**Algorithm 1.**
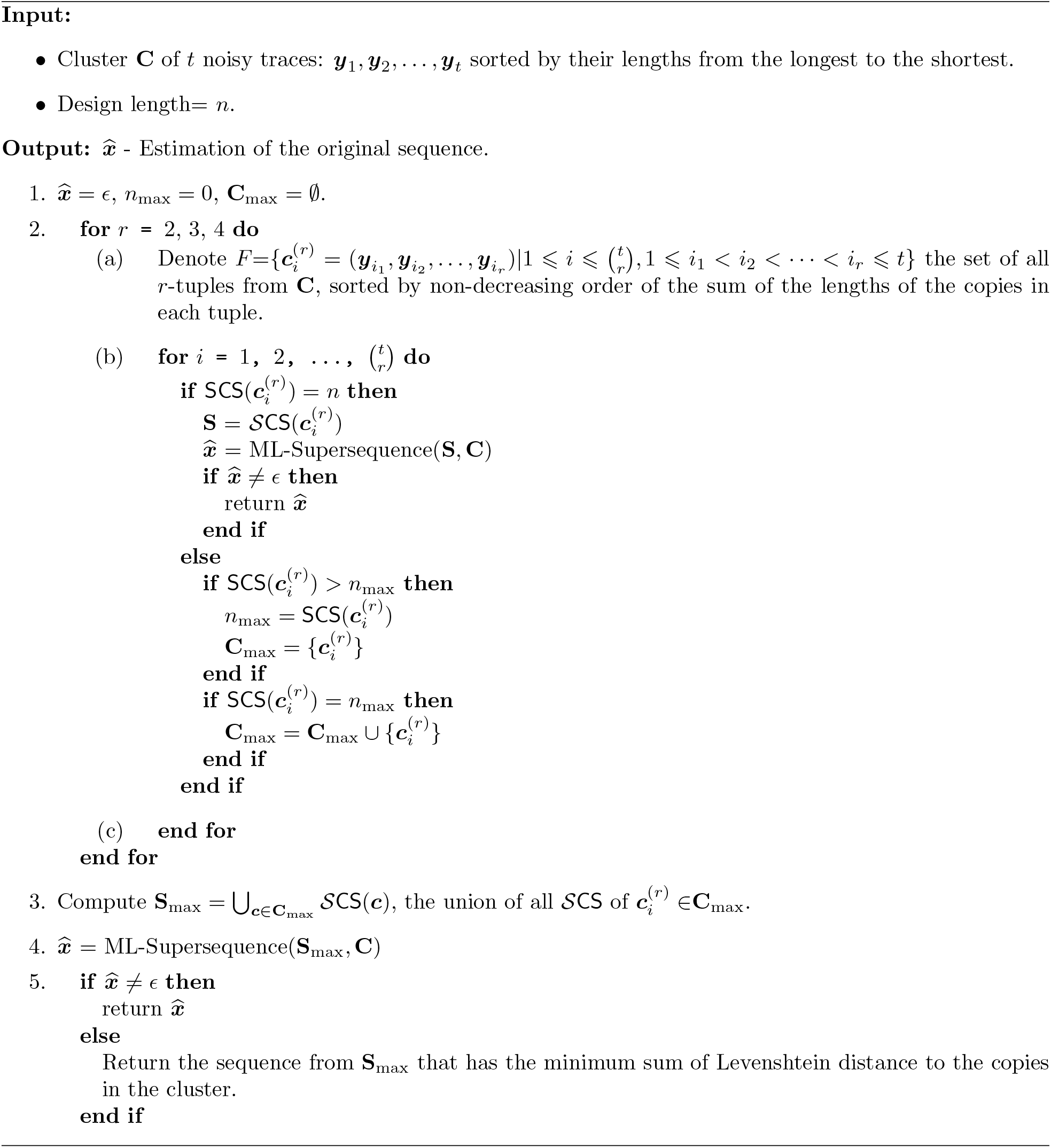
ML-SCS Reconstruction.

### 5.2 Simulations

We evaluated the accuracy and efficiency of Algorithm 1 by the following simulations. These simulations were tested over sequences of length *n* = 200, clusters of size 4 ⩽ *t* ⩽ 10, and deletion probability *p* in the range [0.01, 0.10]. The alphabet size was 4. Each simulation consisted of 100,000 randomly generated clusters. Furthermore, we had another set of simulations for *n* = 100 with deletion probability *p* in the range [0.11, 0.20] and clusters of size 4 ⩽ *t* ⩽ 10. Each simulation for these values of *p,n*, and *t* included 10,000 randomly selected clusters. We calculated the Levenshtein error rate (LER) of the decoded output sequence as well as the average decoding success probability (referred as the *success rate*). We also calculated the *k-error success rate*, which is defined as the fraction of clusters where the Levenshtein distance between the algorithm’s output sequence and the original sequence was at most *k*. Note that for *k* = 0, this is equivalent to calculate the success rate. We also calculated the minimal *k* for which its *k*-error success rate is at least *q*, and denote this value of *k* by *k*_*q* succ_. Note that for *q* = 1 this value determines the minimal number of Levenshtein errors that an error-correcting code must correct in order to fully decode the original sequences using Algorithm 1 with an error-correcting code. In addition, each cluster was also reconstructed using the BMA algorithm [4].

**Algorithm 2.**
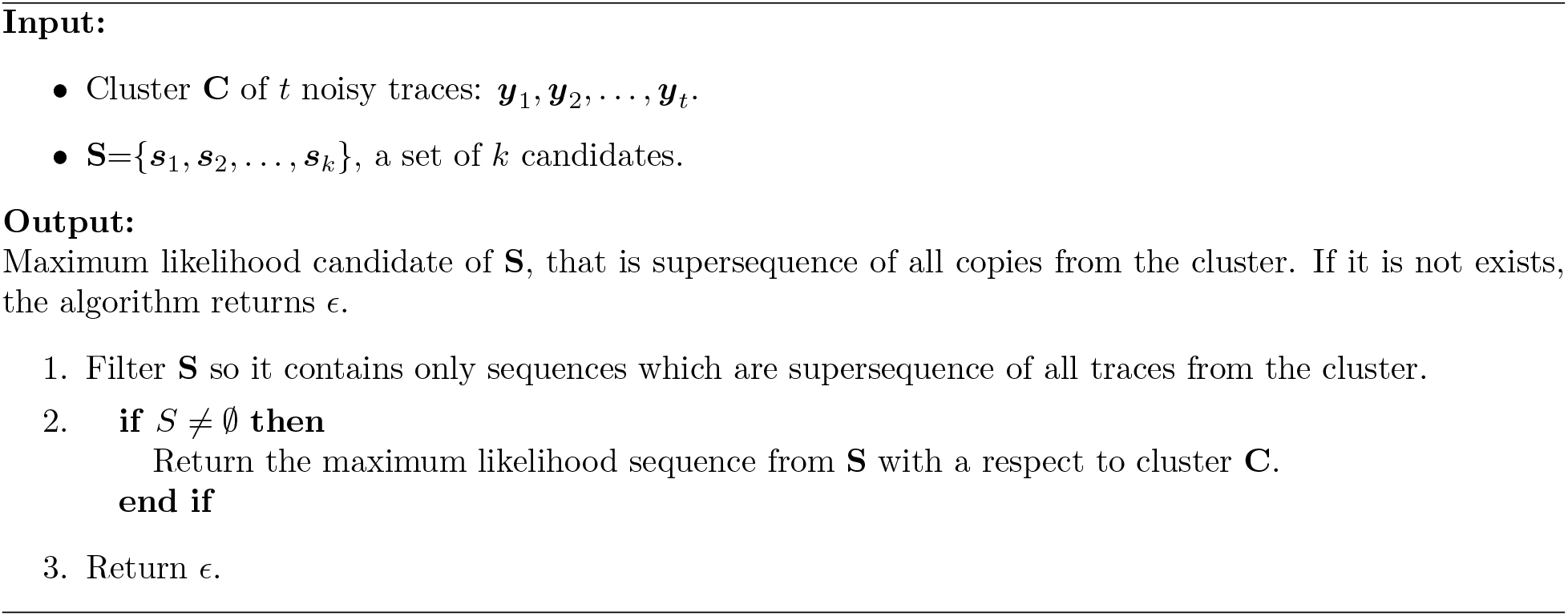
ML-Supersequence.

Figure 1 presents the LER as computed in our simulations of Algorithm 1 and the BMA algorithm for clusters of sizes *t* = 7 and *t* = 10. We also added the trivial lower bound of *p*^*t*^ on the LER [46, 49]. This bound corresponds to the case when the same symbol is deleted in all of the traces. In this case, this symbol will not appear in the list of SCSs of any possible *r*-tuple or even the entire cluster since it cannot be recovered. Hence, it is not possible to recover its value and thus it will be deleted also in the output of the ML decoder.

**Figure 1:**
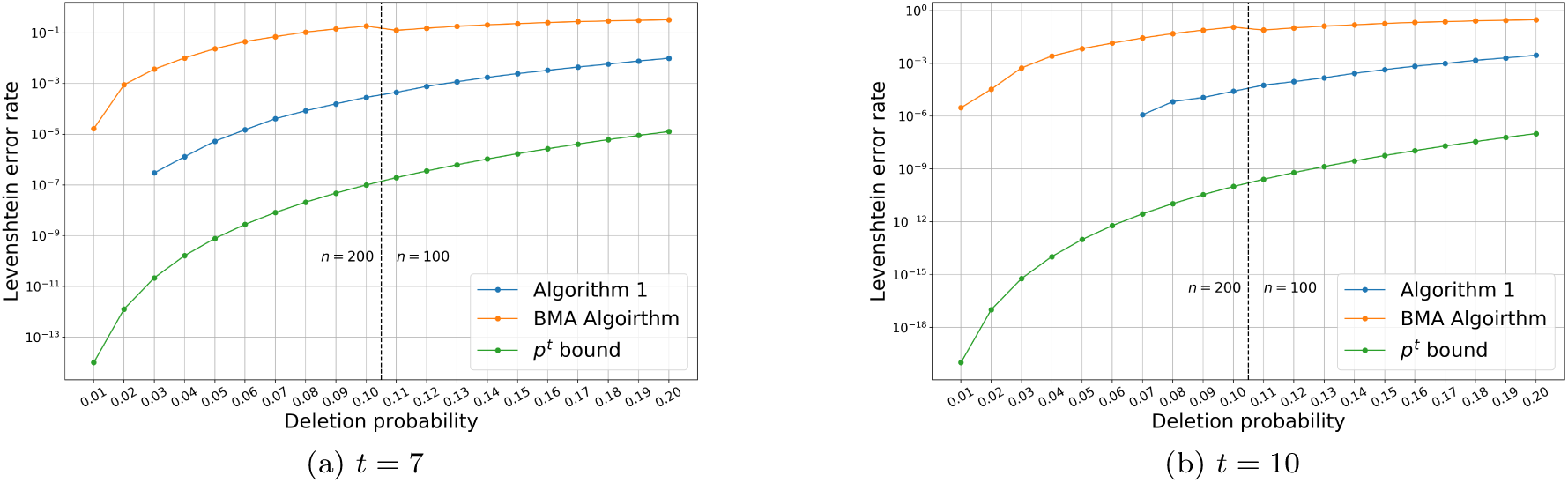
Levenshtein error rate by the deletion probability *p*, for clusters of size 7 (left) and 10 (right). This figure presents results from Algorithm 1, the BMA algorithm [4], and the *p*^*t*^ lower bound. Note that the LER was 0 for *p* ⩽ 0.06 and *p* ⩽ 0.03 for *t* = 10 and *t* = 7, respectively. The *X*-axis represents the different values of the deletion probability in the range [0.01, 0.20] and the *Y* -axis represents the average LER of the clusters.

In order to simulate also high deletion probabilities, we simulated 1000 clusters of sequences over 4-ry alphabet of length *n* = 100 with cluster size *t* between 4 and 10, while the deletion probability was *p* = 0.25. Figure 2(a) presents the *k*-error success rate of this simulation and Figure 2(b) presents the values of *k*_1_succ_ and *k*_0.99_succ_ by the cluster size in the simulation.

**Figure 2:**
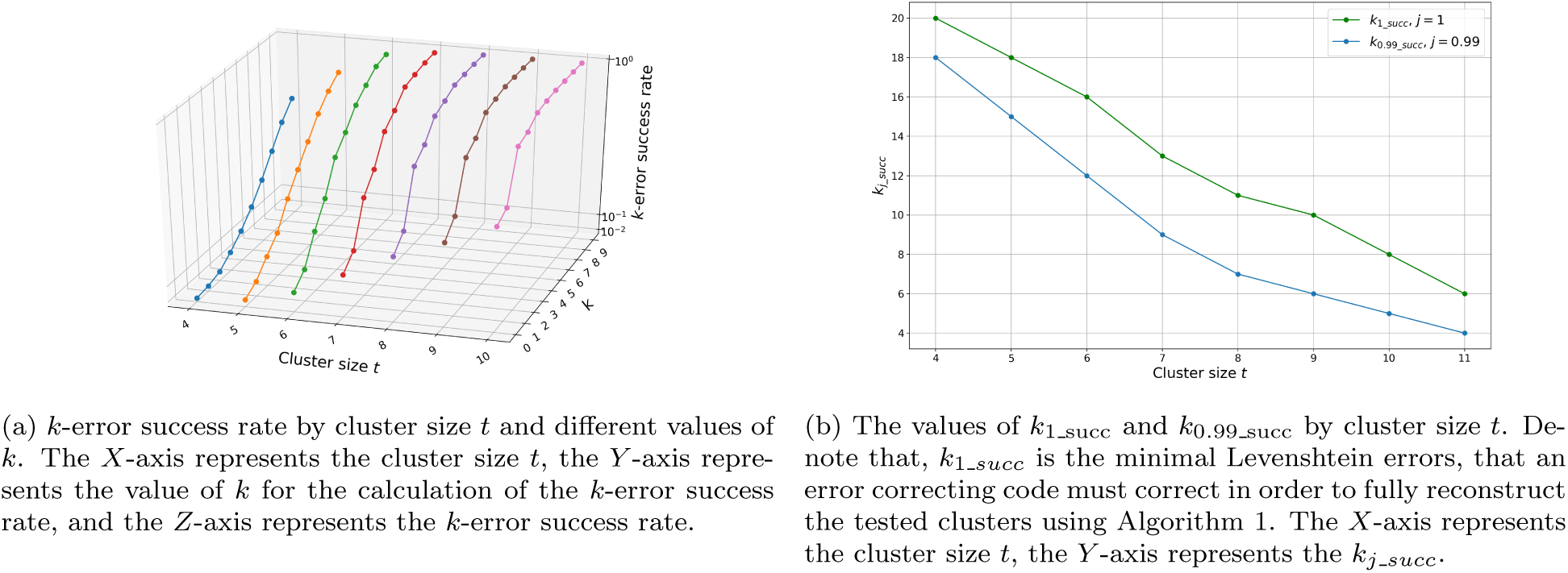
*k*-error success rate, *k*_1_*succ*_ and *k*_*j_succ*_ values by the cluster size 4 ⩽ *t* ⩽ 10. The deletion probability was 0.25.

**Figure 3:**
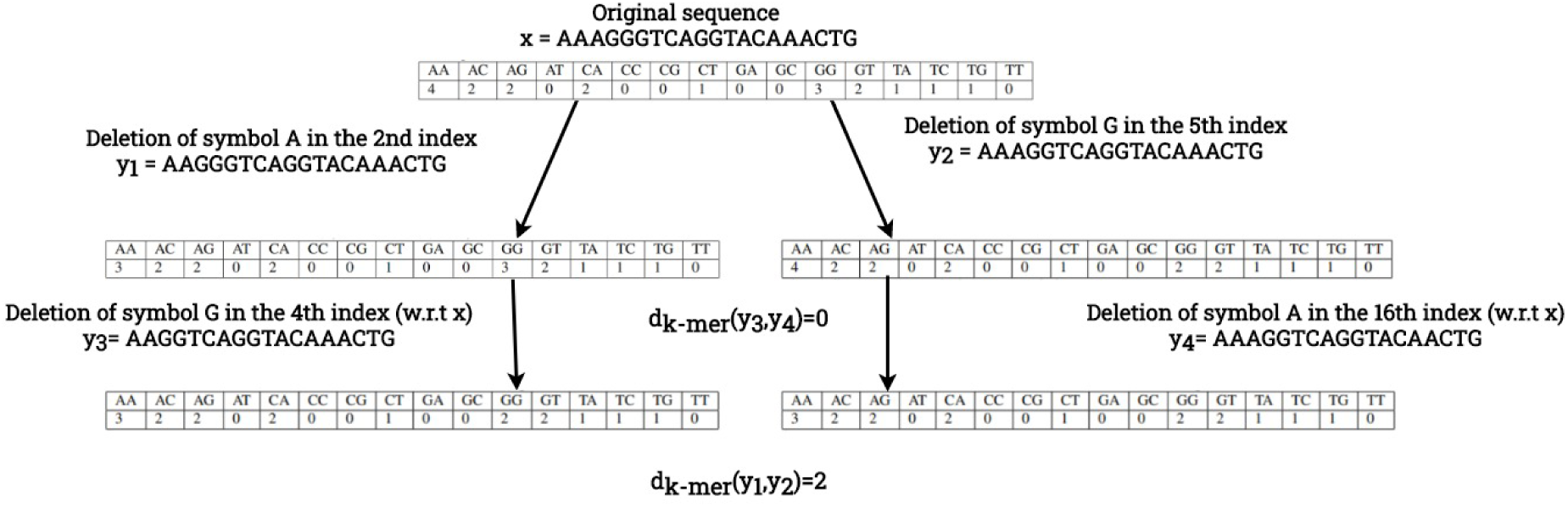
*k*-mer distance demonstration for 4 traces. The original strand ***x*** is of length *n* = 20.

### 5.3 Large Cluster

In case the cluster is of larger size, for example in the order of Θ(*n*), we present in Algorithm 3, a variation of Algorithm 1 for large clusters. In this case, since the cluster is large, the probability to find a pair, triplet, or quadruplet of traces that their set of SCSs contains the original sequence ***x*** is very high, if not even 1. In fact, in all of our simulations, which we will elaborate below in this section, we were always able to successfully decode the original sequence with no errors even when the deletion probability was as high as 0.2. Hence, our main goal in this part is to decrease the runtime of Algorithm 1 while preserving the success rate to be 1. Algorithm 3 keeps the same structure of Algorithm 1, however, it performs two filters on the cluster in order to reduce the computation time.

The complexity of finding the length of the SCS of some set of *r* traces is the multiplication of their lengths, i.e., Θ(*n*^*r*^) [28]. Therefore, the complexity of finding the length of the SCS of a pair of traces is Θ(*n*^2^), while there are Θ(*n*^2^) pairs of traces (assuming the cluster size is Θ(*n*)). Therefore, in this case, calculating the length of the SCS of each pair of traces before considering some triplets is not necessarily the right strategy when our goal is to optimize the algorithm’s running time. Hence, in Algorithm 3 we focused on filtering the traces in the cluster in order to check only a subset of the traces which are more likely to succeed and produce the correct sequence.

To define the filtering criteria for Algorithm 3, we simulated Algorithm 1 on large clusters. The length of the original sequence ***x*** was *n* = 200 and the cluster size was 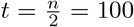. We generated 10,000 clusters of size *t*, where the deletion probability *p* was in the range [0.01, 0.15]. The success rate of all the simulations was 1. We evaluated the percentage of clusters that the first *r*-tuple to have an SCS of length *n* was consisted of the longest 20% traces in the cluster. We observed that when the deletion probability was at most 0.07, in all of the clusters the first *r*-tuple of traces that had an SCS of size *n* consisted from the longest 20% traces in the cluster. For deletion probabilities between 0.08 and 0.11 these percentages ranged between 94.76% and 99.98%, while for *p* = 0.15 this percentage was 60.88%. Therefore, by filtering the longest 20% traces, it was enough to check only 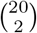 pairs instead of 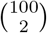 pairs in order to succeed and still reach the successful pair. The results of these simulations are depicted in Figure 4(a).

**Figure 4:**
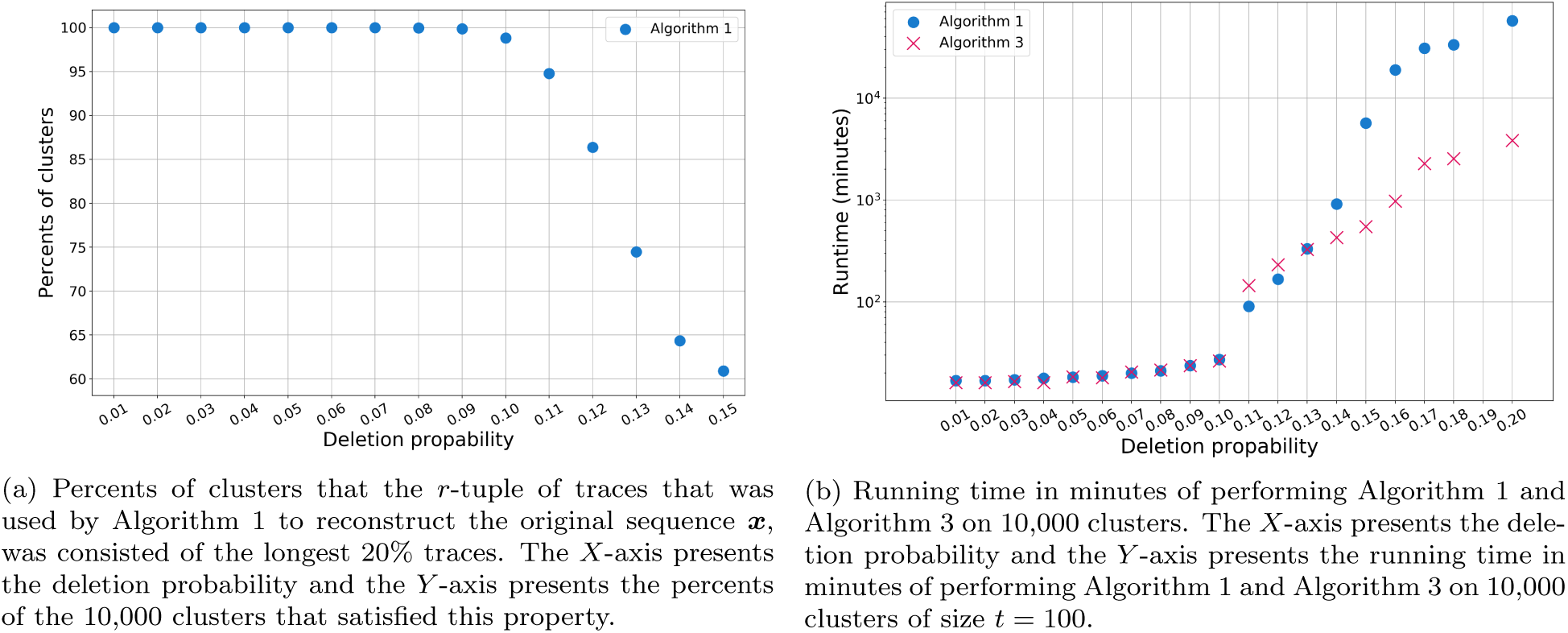
Performance evaluation of Algorithm 1 and Algorithm 3. The simulation were for clusters of size *t* = 100, design length of *n* = 200, for each probability *p* we simulate 10,000 clusters. In all of the simulations the original sequence was reconstructed by the algorithms, introducing a success rate of 1

This observation lead us to the first filter in Algorithm 3, where we picked the longest 20% traces of the cluster. The second filter computes a cost function (in linear time complexity), to be explained below, on a given *r*-tuple of traces in order to evaluate if the traces in this *r*-tuple are likely to have an SCS of length *n*. Thus, the algorithm skips on the SCS computation of *r*-tuples that are less likely to have an SCS of length *n*. First, before performing the first filter, the algorithm calculates the average length of the traces in the cluster and uses it to estimate the deletion probability *p*. Then, if *p >* 0.1, the algorithm calculates the cost function on every *r*-tuple and checks if it is higher than some fixed threshold. This threshold depends on the estimated value of *p* and the cost function is based on a characterization of the sequences, as will be described in Section 5.3.2.

#### 5.3.1 An Algorithm for Large Values of *t*

In this section we present Algorithm 3. We list here the steps that are different from Algorithm 1. In Step 2 the algorithm estimates the deletion probability in the cluster by checking the average length of the traces *n*^*′*^ and then calculates 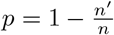. In Step 3, the algorithm filters the cluster so it contains only the longest 20% traces. The last difference between Algorithm 3 and Algorithm 1 can be found in Step 4-b. In this step, before the computation of the SCS of a given *r*-tuple of traces, the algorithm computes the *k*-mer cost function (for *k*-mers of size *k* = 2) and checks if it is larger than the threshold *T*_*p*_.

We evaluated the performance of Algorithm 3 and verified our filters by simulations. Each simulation consisted of 10,000 clusters of size *t* = 100, the length of the original strand was *n* = 200, the alphabet size was *q* = 4, and the deletion probability *p* was in the range [0.01, 0.2]. Algorithm 3 reconstructed the exact sequence ***x*** in all of the tested clusters. A comparison between the runtime of Algorithm 1 and Algorithm 3 can be found in Figure 4(b). Note that we did not compare the running time with the BMA algorithm since its success rate was significantly lower, for example when the deletion probability was 15%, its success rate was roughly 0.46.

#### 5.3.2 The *k*-mer Distance and the *k*-mer Cost Function

The *k-mer vector* of a sequence ***y***, denoted by *k*-mer(***y***), is a vector that counts the frequency in ***y*** of each subsequence of length *k* (*k*-mer). The frequencies are ordered in a lexicographical order of their corresponding *k*-mers. For example for a given sequence ***y*** = “*ACCTCC*” and *k* = 2, its *k*-mer vector is *k*-mer(***y***) = 0100020100000101, according to the following calculation of the frequencies {*AA* : 0, *AC* : 1, *AG* : 0, *AT* : 0, *CA* : 0, *CC* : 2, *CG* : 0, *CT* : 1, *GA* : 0, *GC* : 0, *GG* : 0, *GT* : 0, *T A* : 0, *T C* : 1, *T G* : 0, *T T* : 1}. We define the *k-mer distance* between two sequences ***y***_1_ and ***y***_2_ as the *L*_1_ distance between their *k*-mer vectors. The *k*-mer distance is denoted by *d*_*k*-mer_(***y***_1_, ***y***_2_).

**Algorithm 3.**
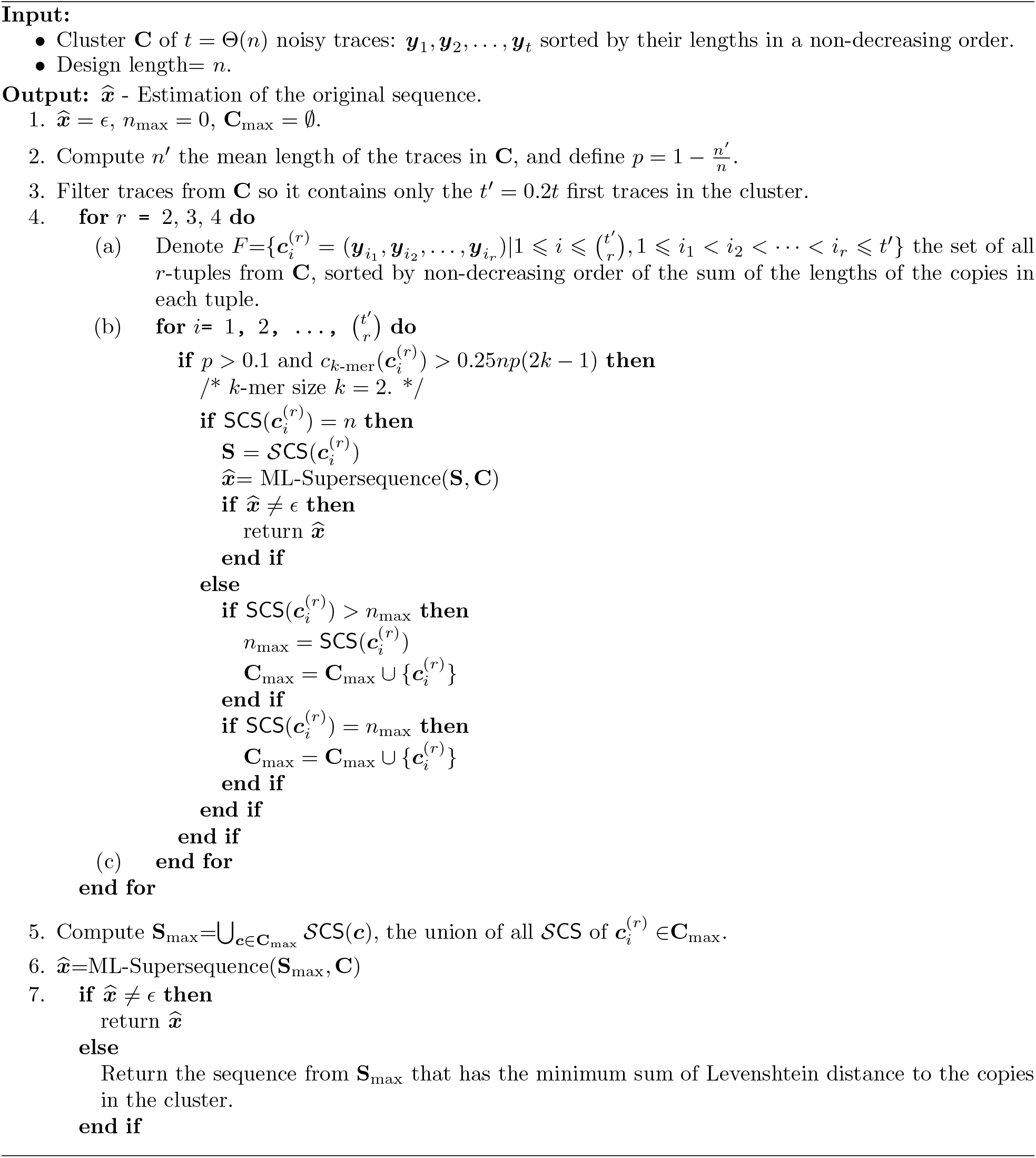
ML-SCS Reconstruction for Large Clusters.

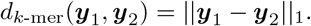

For a given set of *r* sequences **Y** = {****y***_1_, ***y***_2_, …, ***y***_*r*_*}, we define its *k-mer cost function*, which is denoted by *c*_*k*-mer_(***y***_1_, ***y***_2_, …, ***y***_*r*_), as the sum of the *k*-mer distance of each pair of sequences in **Y**. That is,

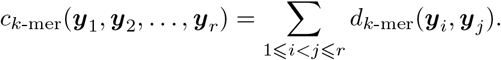

Observe that the *k*-mer distance between a sequence ***x*** and a trace ***y***_1_ which results from ***x*** by one deletion is at most 2*k −* 1. Every deleted symbol in ***x*** decreases the value of at most *k* entries in *k*-mer(***x***) and increases the number of at most *k −* 1 of the entries. Hence, each deletion increases the *k*-mer distance by at most 2*k −* 1, which means that an upper bound on the *k*-mer distance between the original strand ***x*** and a trace ***y***_*i*_ with *np* deletions is *np*(2*k −* 1). However, when comparing the *k*-mer distance of two traces, ***y***_1_ and ***y***_2_, with more than one deletion, the *k*-mer distance can also decrease. An example of such a case is depicted in Figure 3. Combining these two observations, Algorithm 3 estimates if two traces have relatively large Levenshtein distance. If these traces have large Levenshtein distance, it is more likely that both of them will have an SCS of length *n*. Hence, the algorithm checks if the *k*-mer distance is larger than the threshold *T*_*p*_ = 0.25*np*(2*k −* 1) and continues to compute the SCS, only if the condition holds. A similar computation is done for tuples with more than two traces. We use the value of 0.25 in the threshold to consider the cases where the *k*-mer distance decreases as depicted in Figure 3. We selected this value after simulating other values as well, reaching the best result with 0.25. An optimization of this value can be done in further research.

## 6 The DNA Reconstruction Problem

This section studies the DNA reconstruction problem. Assume that a cluster consists of *t* traces, ***y***_1_, ***y***_2_, …, ***y***_*t*_, where all of them are noisy copies of a synthesized strand. This model assumes that every trace is a sequence that is independently received by the transmission of a length-*n* sequence ***x*** (the synthesized strand) through a deletion-insertion-substitution channel with some fixed probability *p*_*d*_ for deletion, *p*_*i*_ for insertion, and *p*_*s*_ for substitution. Our goal is to propose an efficient algorithm which returns 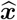, an estimation of the transmitted sequence ***x***, with the intention of minimizing the edit distance between ***x*** and 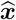. In our simulations, we consider several values of *t* and a wide range of error probabilities as well as data from previous DNA storage experiments.

Before we present the algorithms, we list here several more notations and definitions. An *error vector* of ***y*** and ***x***, denoted by *EV* (***y, x***), is a vector of minimum number of edit operations to transform ***y*** to ***x***. Each entry in *EV* (***y, x***) consists of the index in ***y***, the original symbol in this index, the edit operation and in case the operation is an insertion, substitution the entry also includes the inserted, substituted symbol, respectively. Note that for two sequences ***y*** and ***x***, there could be more than one sequence of edit operations to transform ***y*** to ***x***. The edit distance between a pair of sequences is computed using a dynamic programming table and the error vector is computed by backtracking on this table. Hence, *EV* (***y, x***) is not unique and can be defined uniquely by giving priorities to the different operation in case of ambiguity. That is, if there is an entry in the vector *EV* (***y, x***) (from the last entry to the first), where more than one edit operation can be selected, then, the operation is selected according to these given priorities. The error vector *EV* (***y, x***) also maps each symbol in ***y*** to a symbol in ***x*** (and vice versa). We denote this mapping as *V*_*EV*_ (***y, x***) : {*1, 2, *…*, |***y***|*} →{*1, 2, *…*, |***x***|*} ∪ {*?*}, where *V*_*EV*_ (***y, x***)(*i*) = *j* if and only if the *i*-th symbol in ***y*** appears as the *j*-th symbol in ***x***, with respect to the error vector *EV* (***y, x***). Note that in the case where the *i*-th symbol in ***y*** was classified as a deleted symbol in *EV* (***y, x***), *V*_*EV*_ (***y, x***)(*i*) =?. This mapping can also be represented as a vector of size |***y***|, where the *i*-th entry in this vector is *V*_*EV*_ (***y, x***)(*i*). The *reversed cluster* of a cluster **C**, denoted by **C**^*R*^, consists of the traces in **C** where each one of them is reversed.

### 6.1 The LCS-Anchor Algorithm

In this section we present Algorithm 4, the *LCS-anchor algorithm*. The algorithm receives **C**, a cluster of traces sorted by their lengths, from closest to *n* to the farthest to *n*. First, the algorithm initializes 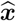 and 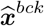 as a length-*n* sequence of the symbol ‘-’. Second, the algorithm computes ***lcs***, an arbitary LCS of ***y***_1_ and ***y***_2_, the two traces in the cluster which their length is closest to *n*. Then, for each of the *t* traces in the cluster, ***y***_*k*_, the algorithm computes *EV* (***y***_*k*_, ***lcs***), and the mapping vector *V*_*EV*_ (***lcs, y***_*k*_). For 1 ⩽ *i* ⩽ |***lcs***| the algorithm performs a majority vote on the *i*-th entries of the *t* mapping vectors. If the majority is *j* ≠ ?, the algorithm writes the symbol ***lcs***(*i*) in the *j*-th index of 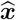. If *j* =? the symbol ***lcs***(*i*) is omitted, and it is not written in 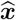. At this point, Algorithm 4 wrote into 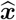 at most LCS(***y***_1_, ***y***_2_) symbols, these symbol serves as “anchor” symbols in the estimated string. Each of the anchor symbols is located in a specific index of 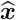, and the rest of the symbols in 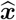 are ‘-’. Note that the anchor symbols are not necessarily placed in consecutive indices of 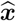. In the following steps, Algorithm 4 computes for all ***y***_*k*_ *∈* **C**, the vectors 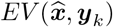 and 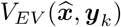. Then, for each *h*, an entry of ‘-’ in 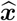, the algorithm performs a majority vote on 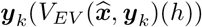 find the most frequent symbol in this entry, and saves it in the *h*-th entry of 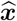, Lastly, the algorithm performs these steps on *C*^*R*^ and saves the resulted sequence in 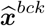. Algorithm 4 returns a output 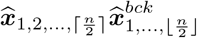.

**Algorithm 4.**
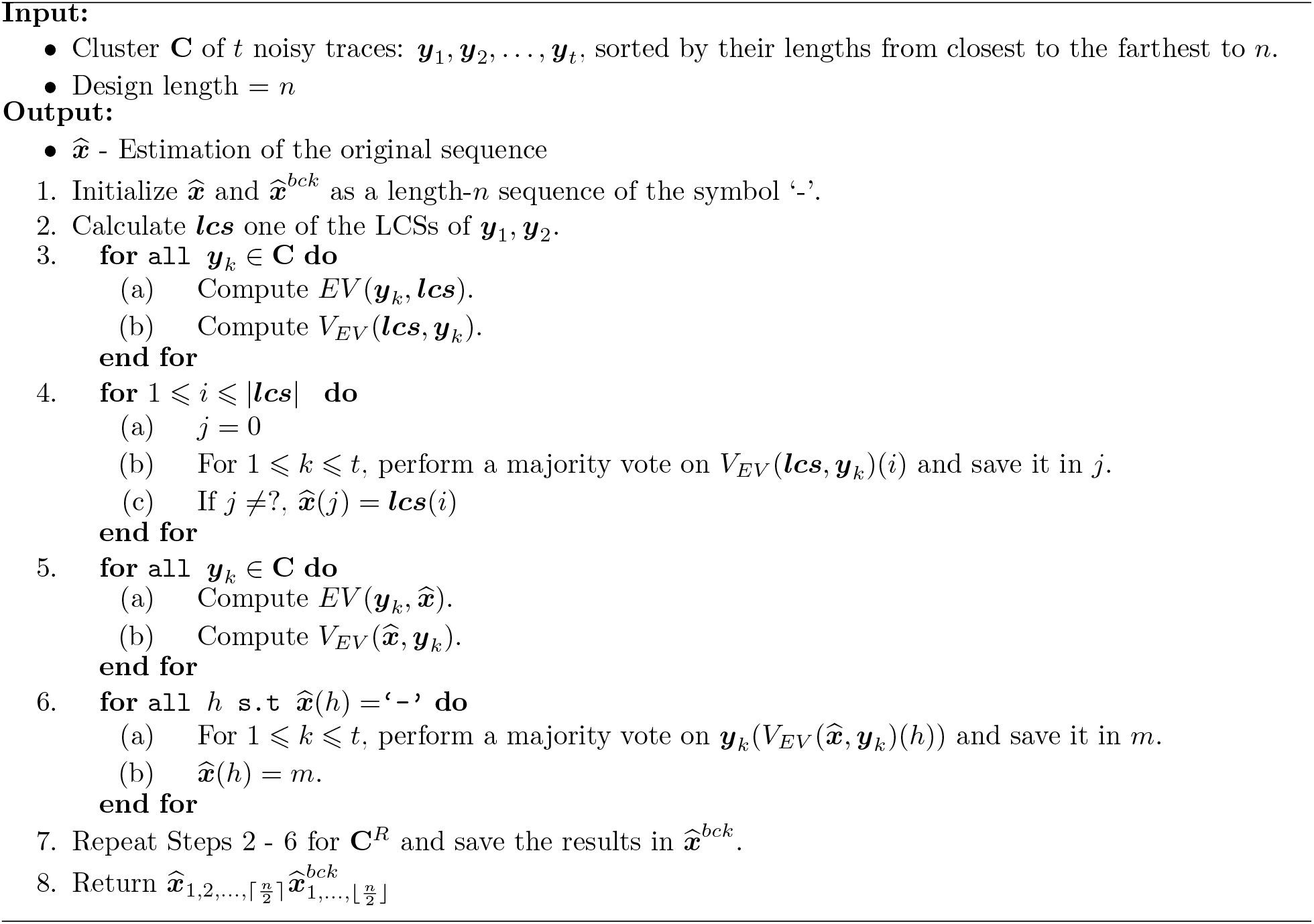
LCS-Anchor.

### 6.2 The Iterative Reconstruction Algorithm

In this section we present Algorithm 5. The algorithm receives a cluster of *t* traces **C** and the design length *n*. Algorithm 5 uses several methods to revise the traces from the cluster and to generate from the revised traces a multiset of candidates. Then, Algorithm 5 returns the candidate that is most likely to be the original sequence ***x***. The methods used to revise the traces are described in this section as Algorithm 6 and Algorithm 7. Algorithm 5 invokes Algorithm 6 and Algorithm 7 on the cluster in two different procedures as described in Algorithm 8 and Algorithm 9.

The first method is described in Algorithm 6. The algorithm receives **C**, a cluster of *t* traces, the design length *n*, and ***y***_*i*_, a trace from the cluster. Algorithm 6 calculates for every 1 ⩽ *k* ⩽ *t, k ≠ i*, the vector *EV* (***y***_*i*_, ***y***_*k*_). In some fo the cases, there may be more than one error vector for *EV* (***y***_*i*_, ***y***_*k*_), which corresponds to the edit operations to transform ***y***_*i*_ to ***y***_*k*_. In these cases, the algorithm prioritizes substitutions, then insertions, then deletions in order to choose one unique vector^2^. Then, the algorithm performs a majority vote in each index on these vectors and creates **S**, which is a vector of edit operations. Lastly, Algorithm 6 performs the edit operations on ***y***_*i*_, and returns it as an output for Algorithm 8 and Algorithm 9. Algorithm 6 is used as a procedure in Algorithm 8 and Algorithm 9 to correct substitution and insertion errors of the traces in the cluster.

The second method is described in Algorithm 7. Similarly to Algorithm 6, Algorithm 7 receives **C**, a cluster of *t* traces, the design length *n*, and ***y***_*k*_, a trace from the cluster. Algorithm 7 uses similar patterns (defined in Section 6.2.1) on each pair of traces and creates a weighted graph from them. Each vertex of the graph represents a pattern, and an edge connects patterns with identical prefix and suffix. The weight on each edge represents the frequency of the incoming pattern, the number of pairs of traces in the cluster that have this pattern in their sequences. Algorithm 7 is described in detail in Section 6.2.1. Algorithm 7 is used as a procedure in in Algorithm 8 and Algorithm 9 to correct deletion errors in the traces in the cluster.

Algorithm 8 receives a cluster of *t* traces **C** and the design length *n*. Algorithm 8 performs *k* cycles, where in each cycle it iterates over all the traces in the cluster. For each trace ***y***_*k*_, it first uses Algorithm 6 to correct substitution errors, then it uses Algorithm 7 to correct deletion errors, and lastly, it uses Algorithm 6 to correct insertion errors. When it finishes iterating over the traces in the cluster, Algorithm 8 updates the cluster with all the revised traces and continues to the next cycle. At the end, Algorithm 8 performs the same procedure on **C**^*R*^. Algorithm 8 returns a multiset of all the revised traces.

Algorithm 9 also receives a cluster of *t* traces **C** and the design length *n*. Algorithm 9 uses the same procedures as Algorithm 8. However, in each cycle, it first corrects substitutions in all of the traces in the cluster using algorithm 6, then it invokes algorithm 7 on each trace to correct deletions, and finally invokes Algorithm 6 to correct insertions. Similarly to Algorithm 8, Algorithm 9 performs the same operations also on **C**^*R*^ and returns a multiset of the results.

Algorithm 5 invokes Algorithms 8 and 9, with *k* = 2 cycles and combines the resulted multisets to the multiset **S**. If one or more sequences of length *n* exists in the multiset **S**, it returns the one that minimizes the sum of edit distances to the traces in the cluster. Otherwise, it checks if there are sequences of length *n −* 1 or *n* + 1 in **S**, and returns the most frequent among them. If such a sequence does not exists, it returns the first sequence in **S**.

**Algorithm 5.**
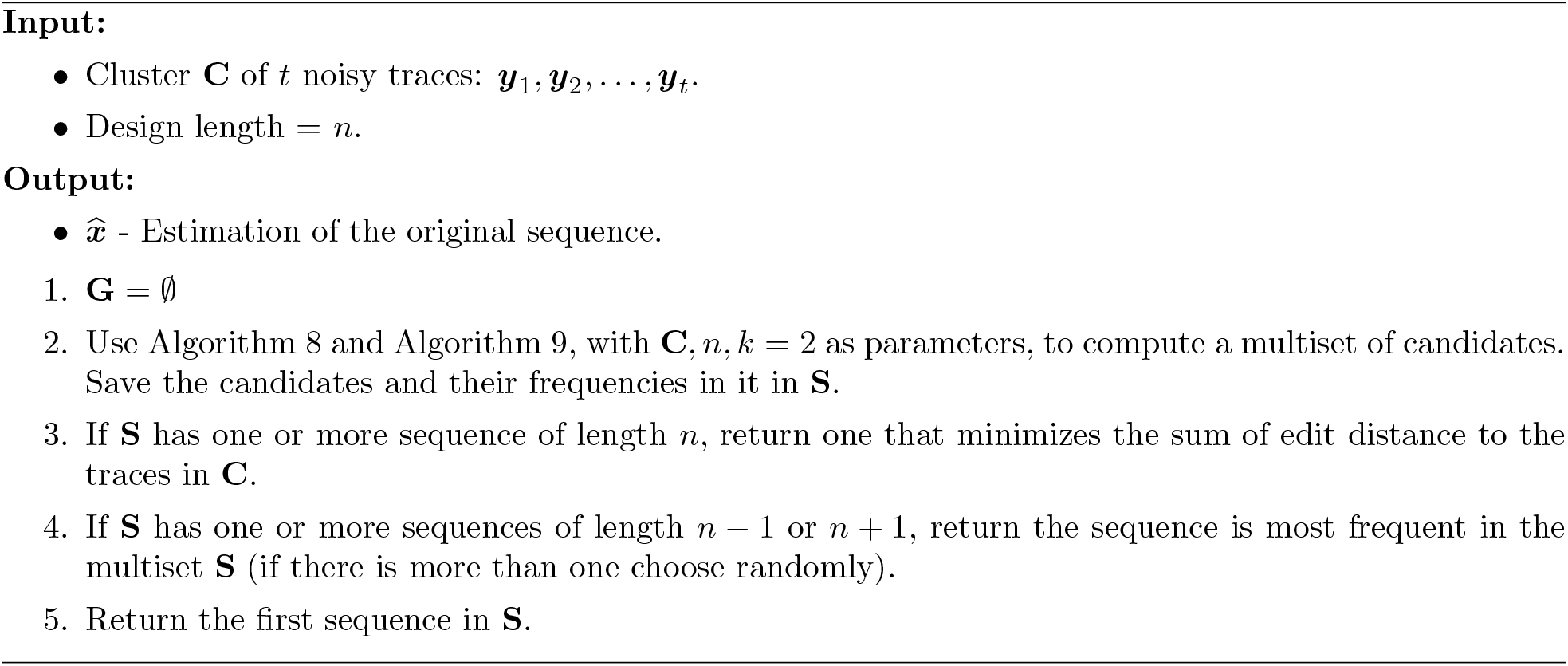
Iterative Reconstruction.

#### 6.2.1 The Pattern Path Algorithm

In this section we present Algorithm 7, the Pattern-Path algorithm. Algorithm 7 is being used to correct deletion errors. Denote by ***w*** an arbitrary LCS sequence of ***x*** and ***y*** of length ℓ. The sequence ***w*** is a subsequence of ***x***, and hence, all of its ℓ symbols appear in some indices of ***x***^3^, and assume these indices are given by 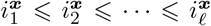. Furthermore, we also define the *embedding sequence* of ***w*** in ***x***, denoted by ***u***_***x***,***w***_, as a sequence of length |***x***| where for 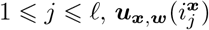 equals to 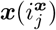 and otherwise it equals to ?.

The *gap* of ***x, y*** and their LCS sequence ***w*** in index 1 ⩽ *j* ⩽ |***x***| with respect to ***u***_***x***,***w***_ and ***u***_***y***,***w***_, denoted by 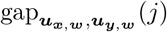, is defined as follows. In case the *j*-th or the (*j −* 1)-th symbol in ***u***_***x***,***w***_ equals ?, 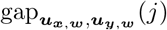 is defined as an empty sequence. Otherwise, the symbol ***u***_***x***,***w***_(*j*) also appears in ***w***. Denote by *j*^*′*^, the index of the symbol ***u***_***x***,***w***_(*j*) in ***w***. The sequence ***w*** is an LCS of ***x*** and ***y***, and ***u***_***y***,***w***_ is the embedding sequence of ***w*** in ***y***. Given ***u***_***y***,***w***_, we can define one set of indices 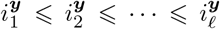 such that 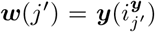 for 1 ⩽ *j* ⩽ ℓ. Given such a set of indices, 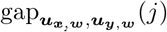 is defined as the sequence 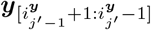, which is the sequence between the appearances of the *j*^*/*^-th and the (*j*^*/*^ *−* 1)-th symbols of ***w*** in ***y***. Note that since 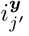 can be equal to 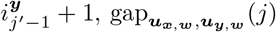 can be an empty sequence.

The *pattern* of ***x*** and ***y*** with respect to the LCS sequence ***w***, its embedding sequences ***u***_***x***,***w***_ and ***u***_***y***,***w***_, an index 1 ⩽ *i* ⩽ |***x***| and a length *m* ⩾ 2, denoted by *Ptn*(***x, y, w, u***_***x***,***w***_, ***u***_***y***,***w***_, *i, m*), is defined as:

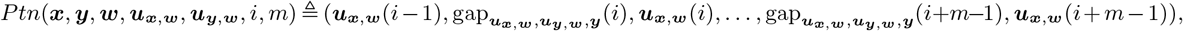

where for *i <* 1 and *i >* |*x*|, the symbol ***u***_***x***,***w***_(*i*) is defined as the null character and 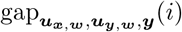 is defined as an empty sequence.

We also define the prefix and suffix of a pattern *Ptn*(***x, y, w, u***_***x***,***w***_, ***u***_***y***,***w***_, *i, m*) to be:

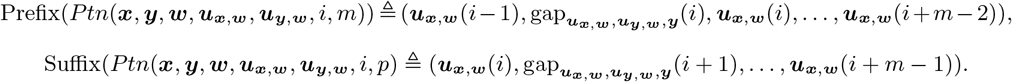

Finally, we define

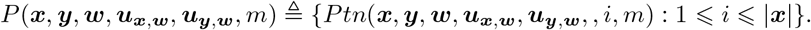

Algorithm 7 receives a cluster **C** of *t* traces and one of the traces in the cluster ***y***_*k*_. First, the algorithm initializes *L*[***y***_*k*_], which is a set of |***y***_*k*_ | empty lists. For 1 ⩽ *i* ⩽|*y*_*k*_|, the *i*-th list of *L*[***y***_*k*_] is denoted by *L*[***y***_*k*_]_*i*_. Algorithm 7 pairs ***y***_*k*_ with each of the other traces in **C**. For each pair of traces, ***y***_*k*_ and ***y***_*h*_, Algorithm 7 computes an arbitrary LCS sequence ***w***, and an arbitrary embedding sequence 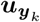. Then it uses ***w*** and 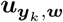 to computes 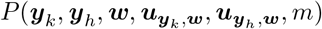. For 1 ⩽ *i* ⩽|*y*_*k*_|, the algorithm saves 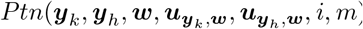 in *L*[***y***_*k*_]_*i*_. Then, Algorithm 7 builds the *pattern graph G*_*pat*_(***y***_*k*_) = (*V* (***y***_*k*_), *E*(***y***_*k*_)), which is a directed acyclic graph, and is defined as follows.

1. 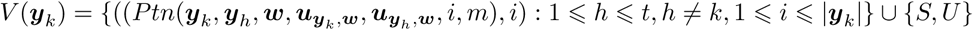, *i*) : 1 ⩽ *h* ⩽ *t, h ≠ k*, 1 ⩽ *i* ⩽ |***y***_*k*_ |*} ∪ {S, U}*. The vertices are pairs of pattern and their index. Note that the same pattern can appear in several vertices with different indices *i*. The value |*V* | equals to the number of distinct pattern-index pairs.
2. 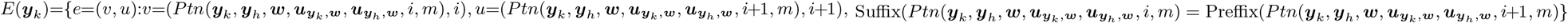.
3. The weights of the edges are defined by *w* : *E → N* as follows: For *e* = (*v, u*), where 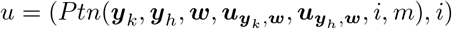, it holds that

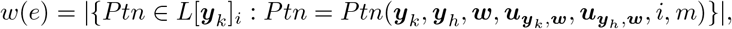

which is the number of appearances of 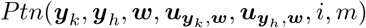.
4. The vertex *S* which does not correspond to any pattern, is connected to all vertices of the first index. The weight of these edges is the number of appearances of the incoming vertex pattern.
5. The vertex *U* has incoming edges from all vertices of the last index and the weight of each edge is zero.

Algorithm 7 finds a longest path from *S* to *U* in the graph. This path induces a sequence, denoted by 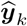, that consists of patterns of ***y***_*k*_ where some of them include gaps. The algorithm returns 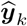, which is a revised version of ***y***_*k*_, while adding to the original sequence of ***y***_*k*_ the gaps that are inherited from the longest path vertices.

**Example 1**. We present here a short example of the definitions above and Algorithm 7. The original strand in this example is ***x*** and the cluster of traces is **C** = ***y***_1_, …, ***y***_5_, Note that the original length is *n* = 10. The traces are noisy copies of ***x*** and include deletions, insertions, and substitutions. In this example Algorithm 7 receives the cluster **C** and the trace ***y***_*k*_ = ***y***_1_ as its input.

- ***x*** = *GTAGTGCCTG*.
- ***y***_1_ = *GTAGGTGCCG*.
- ***y***_2_ = *GTAGTCCTG*.
- ***y***_3_ = *GTAGTGCCTG*.
- ***y***_4_ = *GTAGCGCCAG*.
- ***y***_5_ = *GCATGCTCTG*.

Figure 5 presents the process of computing the patterns of (***y***_1_, ***y***_2_), (***y***_1_, ***y***_3_), (***y***_1_, ***y***_4_), (***y***_1_, ***y***_5_). For each pair, ***y***_1_ and ***y***_*i*_, Figure 5 depicts ***w***_*i*_, which is an LCS of the sequences ***y***_1_ and ***y***_*i*_. Then, the figure presents 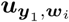 and 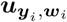, which are the embedding sequences that Algorithm 7 uses in order to compute the patterns. Lastly, the list of patterns of each pair is depicted in an increasing order of their indices. Note that lowercase symbols present gaps and *X* presents the symbol ?. The following list summarizes the patterns and their frequencies. Each list includes patterns from specific index. The numbers on the right side of each pattern in a list represents the pattern’s frequency.

**Figure 5:**
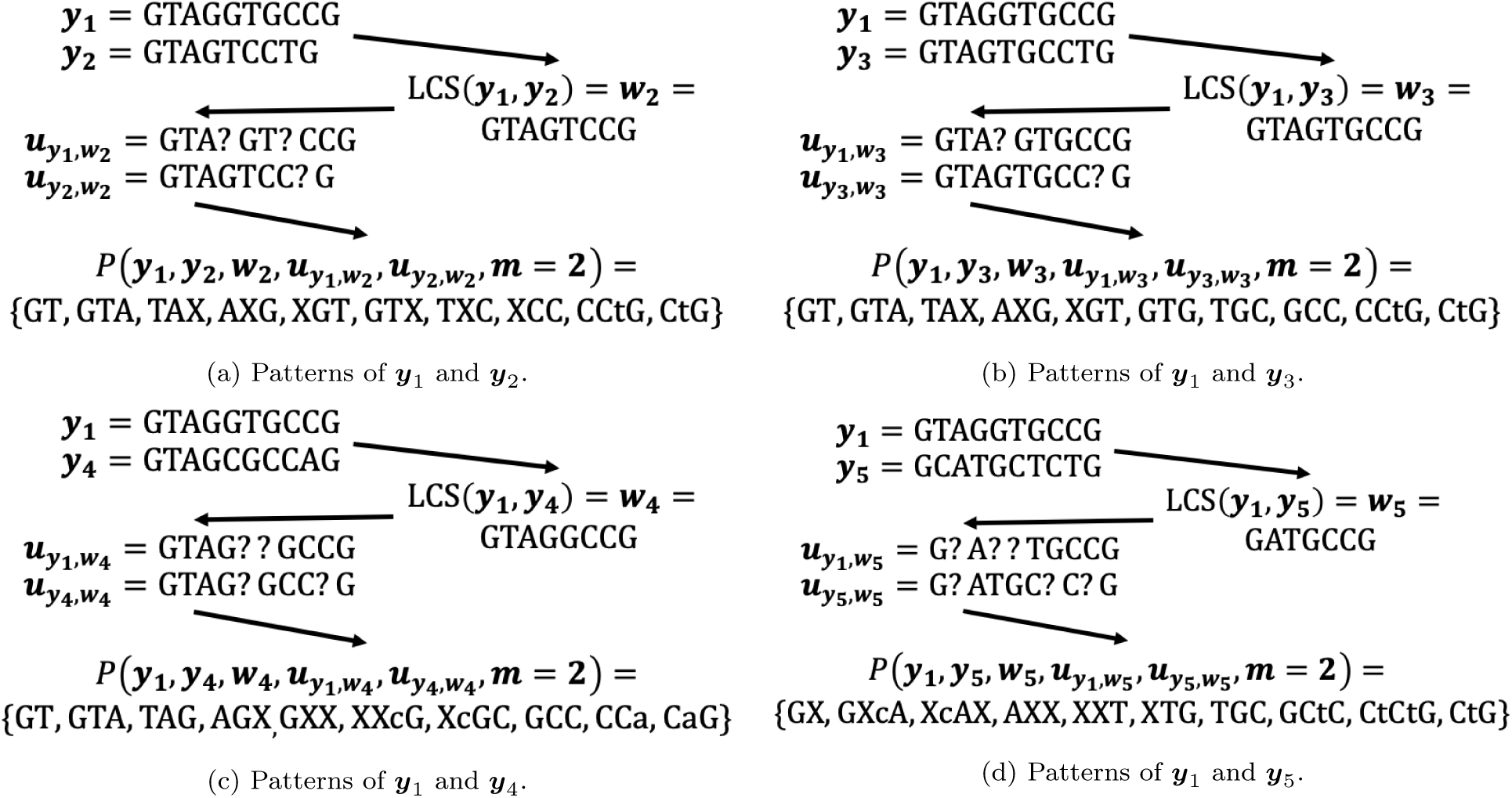
Algorithm 7 Example - Patterns of *y*_1_.

- *L*[***y***_1_]_1_ = {*GT* : 3, *GX* : 1}.
- *L*[***y***_1_]_2_ = {*GTA* : 3, *GXcA* : 1}.
- *L*[***y***_1_]_3_ = {*TAX* : 2, *TAG* : 1, *XcAX* : 1}.
- *L*[***y***_1_]_4_ = {*AXG* : 2, *AGX* : 1, *AXX* : 1}.
- *L*[***y***_1_]_5_ = {*XGT* : 2, *GXX* : 1, *XXT* : 1}.
- *L*[***y***_1_]_6_ = {*GTG* : 1, *GTX* : 1, *XXcG* : 1, *XTG* : 1}.
- *L*[***y***_1_]_7_ = {*TGC* : 2, *TXC* : 1, *XcGC* : 1}.
- *L*[***y***_1_]_8_ = {*GCC* : 2, *SCC* : 1, *GCtG* : 1}.
- *L*[***y***_1_]_9_ = {*CCtG* : 2, *CCa* : 1.*CtCtG* : 1}.
- *L*[***y***_1_]_10_ = {*CtG* : 3, *CaG* : 1}.

It is not hard to observe that the longest path in the pattern path graph of this example is:

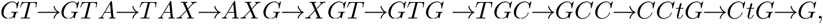

and the algorithm output will be 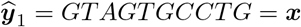.

**Algorithm 6.**
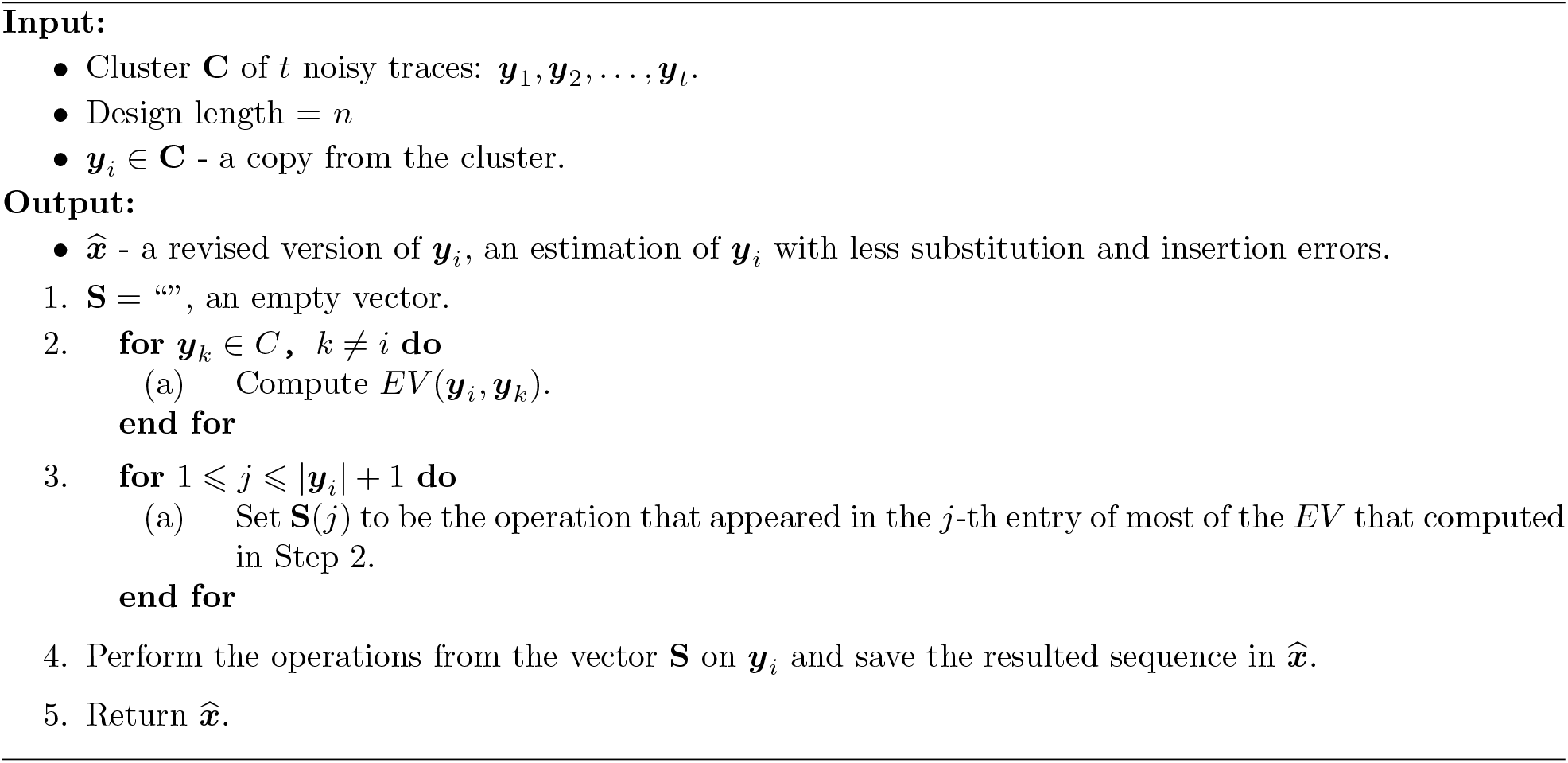
Error Vectors Majority.

**Algorithm 7.**
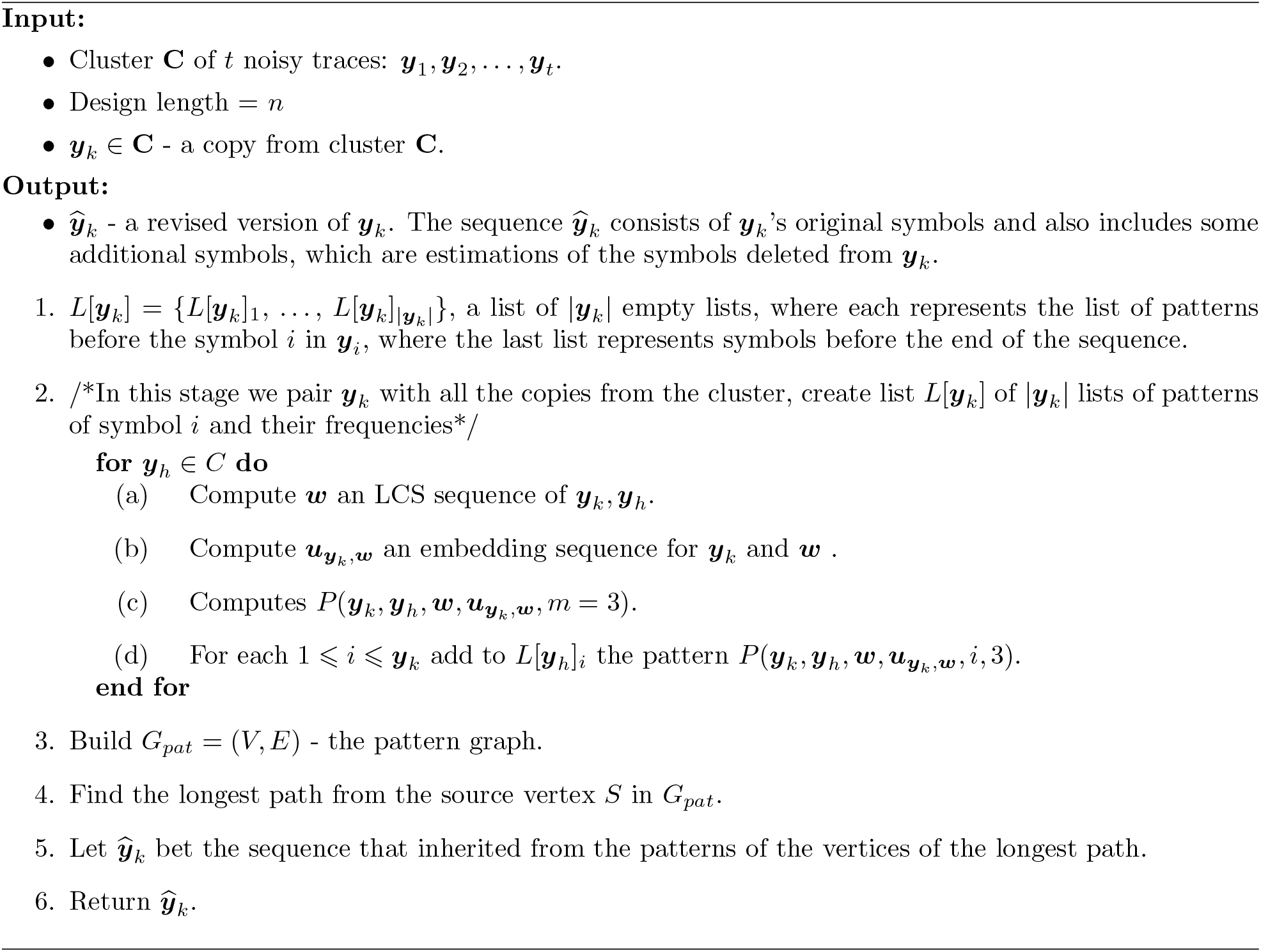
Pattern-Path.

**Algorithm 8.**
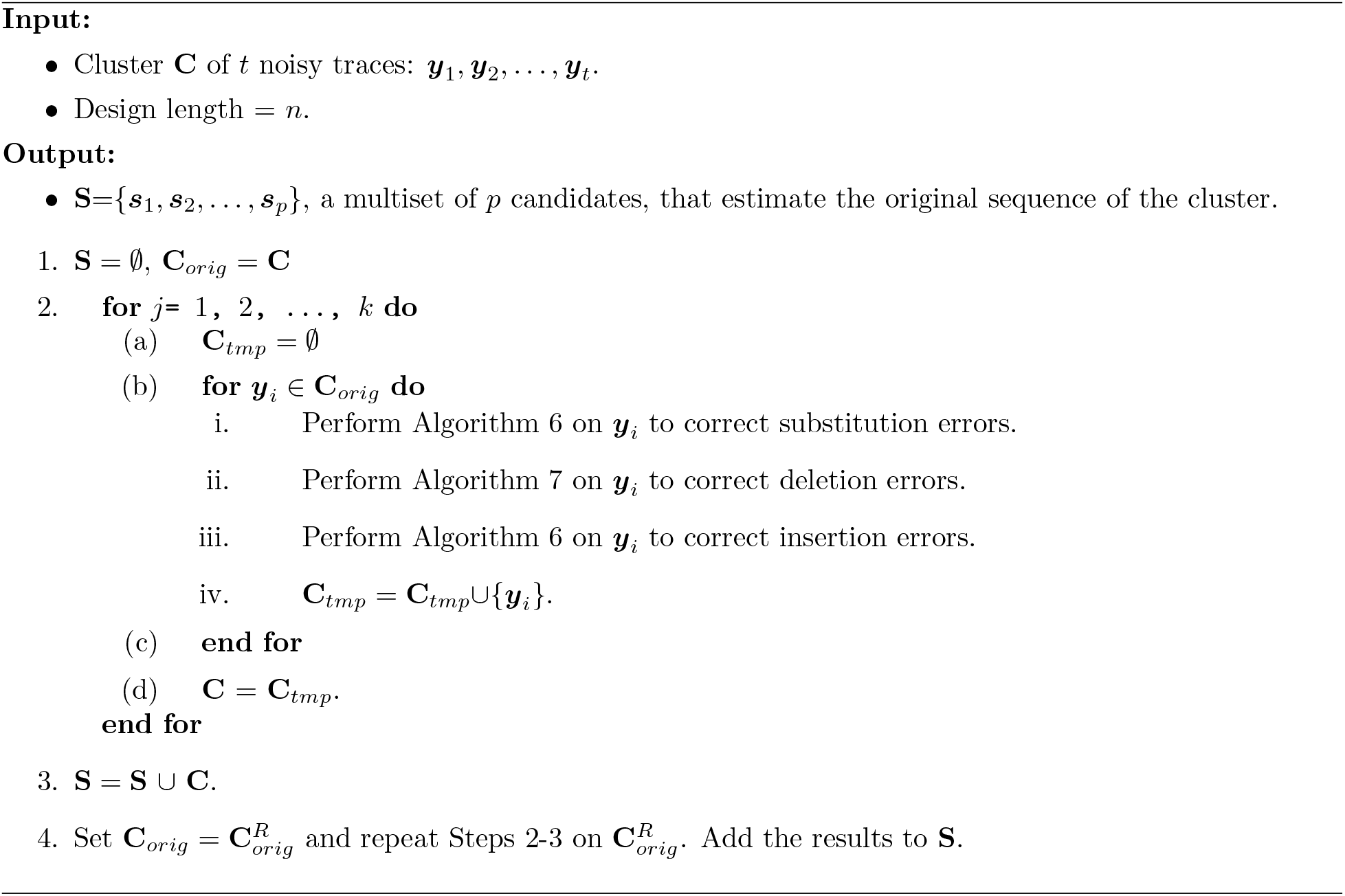
Iterative Reconstruction - Horizontal.

## 7 Results

In this section we present an evaluation of the accuracy of Algorithm 4 and Algorithm 5 on simulated data and on data from previous DNA storage experiments [20, 40, 24]. We also implemented the algorithms from [23]^4^ and from [52]^5^, and our variation of the BMA algorithm [4] to support also insertion and substitution errors, which is referred by the *Divider BMA algorithm*.

The Divider BMA algorithm receives a cluster and the design length *n*. The Divider BMA algorithm divides the traces of the cluster into three sub-clusters by their lengths, traces of length *n*, traces of length smaller than *n* and traces of length larger than *n*. It performs a majority vote on the traces of length *n*. Then, similarly to the technique presented in the BMA algorithm [4] and in [23], the Divider BMA algorithm performs a majority vote on the sub-cluster of traces of length smaller than *n*, while detecting and correcting deletion errors. Lastly, the Divider BMA algorithm uses the same technique on the traces of length larger than *n* to detect and correct insertion errors.

We compare the edit error rates and the success rates of all the algorithms. In all of the simulations, Algorithm 5 presented significantly smaller edit error rates and higher success rates. The results on the simulated data are depicted in Figure 6, Figure 7, and Figure 8. The results on the data from previous DNA storage experiments can be found in Figure 9.

**Figure 6:**
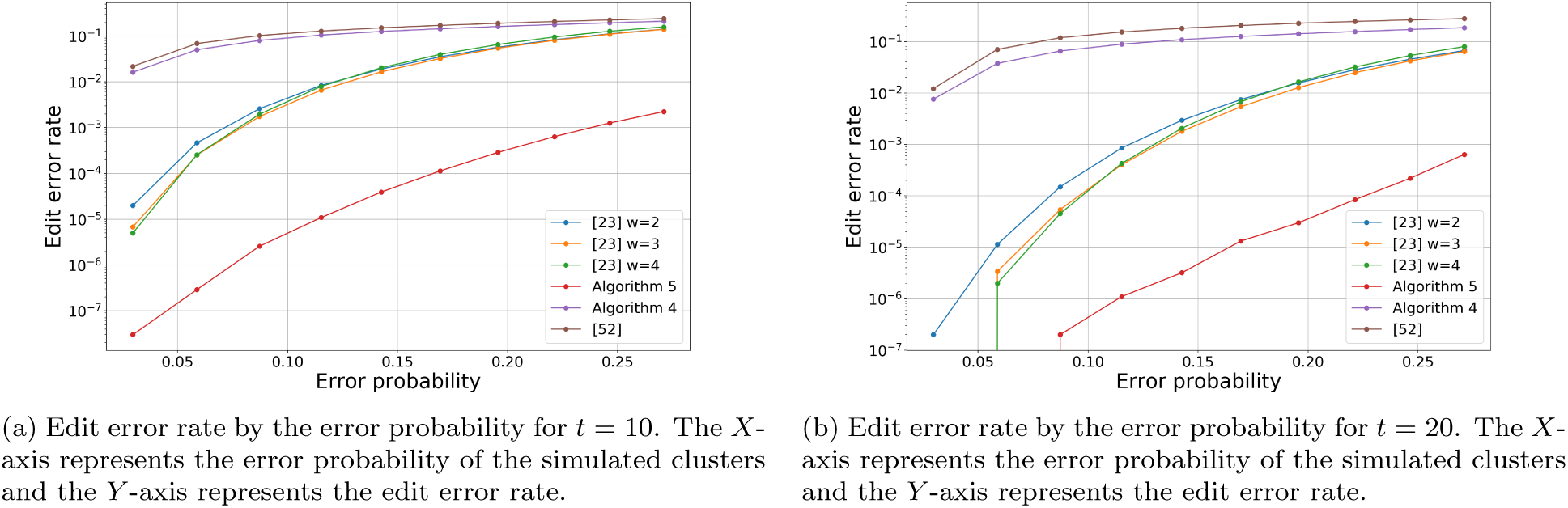
Edit error rate by the error probabilities for the cluster sizes *t* = 10 and *t* = 20. The length of the original sequence was *n* = 100 and the error probabilities ranges between 0.029701 and 0.271.

**Figure 7:**
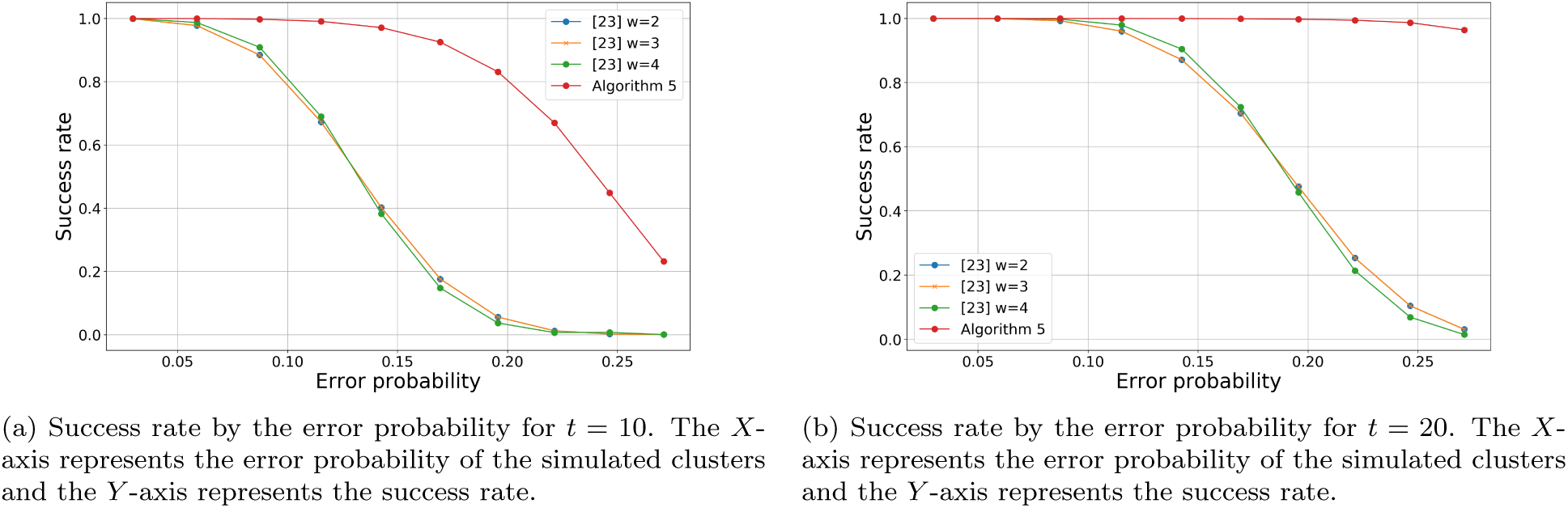
Success rate by the error probabilities for the cluster sizes *t* = 10 and *t* = 20. The length of the original sequence was *n* = 100 and the error probabilities ranges between 0.029701 and 0.271.

**Figure 8:**
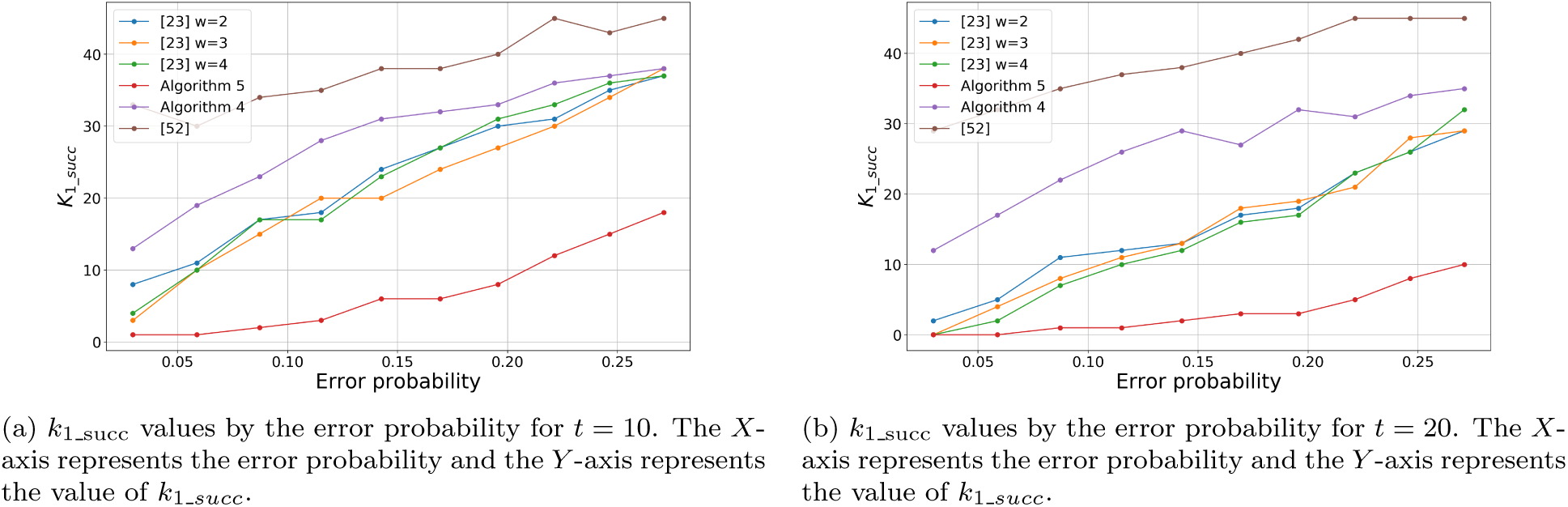
*k*_1_succ_ values by the error probabilities for the cluster sizes *t* = 10 and *t* = 20. The length of the original sequence was *n* = 100 and the error probabilities ranges between 0.029701 and 0.271.

**Figure 9:**
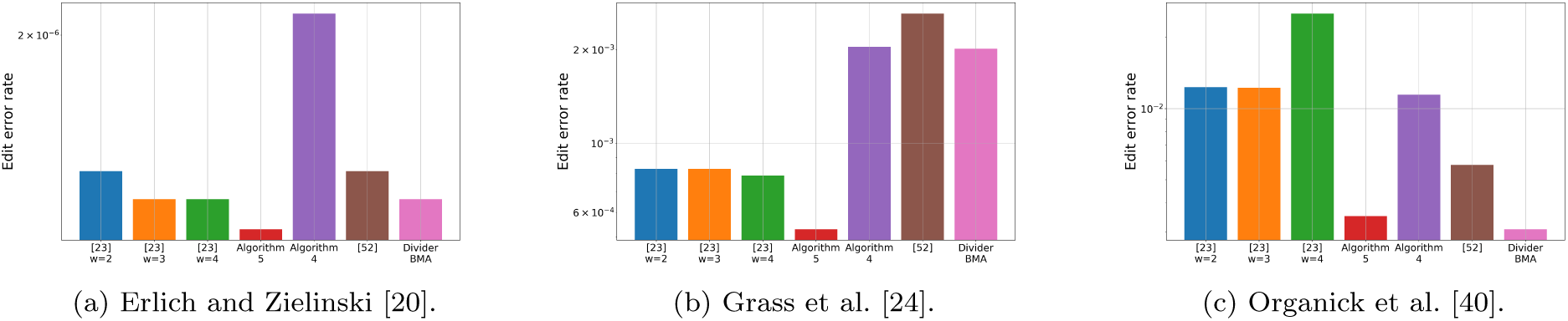
Edit error rate by the reconstruction algorithm, for data from DNA storage experiments [20, 24, 40].

### 7.1 Results on Simulated Data

We evaluated the accuracy of Algorithm 4 and Algorithm 5 by simulations. First, we present our interpretation of the deletion-insertion-substitution channel. In our deletion-insertion-substitution channel, the sequence is transmitted symbol-by-symbol. First, before transmitting the symbol, it checks for an insertion error before the transmitted symbol. The channel flips a coin, and in probability *p*_*i*_, an insertion error occurs before the transmitted symbol. If an insertion error occurs, the inserted symbol is chosen uniformly. Then, the channel checks for a deletion error, and again flips a coin, and in probability *p*_*d*_ the transmitted symbol is deleted. Lastly, the channel checks for a substitution error. The channel flips a coin, and in probability *p*_*s*_ the transmitted symbol is substituted to another symbol. The substituted symbol is chosen uniformly. In case that both deletion and substitution errors occurs in the same symbol, we refer to it as a substitution.

**Algorithm 9.**
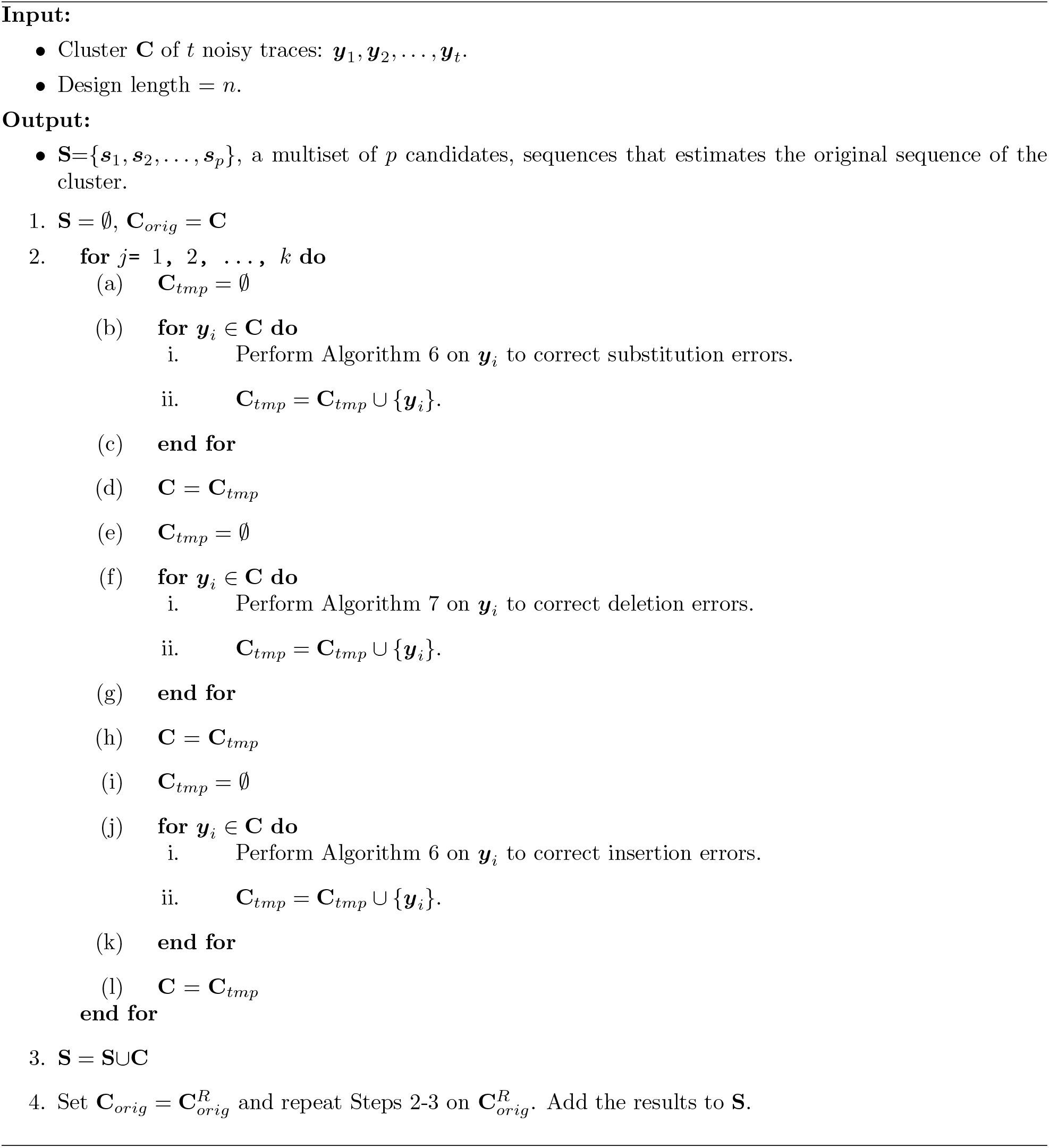
Iterative Reconstruction - Vertical.

We simulated 100,000 clusters of sizes *t* = 6, 10, 20, the sequences length was *n* = 100, and the alphabet size was *q* = 4. The deletion, insertion, and substitution probabilities were all identical, and ranged between 0.01 and 0.1. It means that the actual error probability of each cluster was 1 *−* (1 *− p*_*i*_)(1 *− p*_*s*_)(1 *− p*_*d*_) and ranged between 0.029701 and 0.271. We reconstructed the original sequences of the clusters using Algorithm 4, Algorithm 5 and the algorithms from [23] and from [52]. For each algorithm we evaluated its edit error rate, the success rate, and the value of *k*_1_succ_. The edit error rate of Algorithm 5 was the lowest among the tested algorithms, while the algorithm from [52] presented the highest edit error rates. Moreover, it can be seen that Algorithm 5 presented significantly low edit error rates value for higher values of error probabilities. In addition, Algorithm 5 also presented the lowest value of *k*_1_succ_. For example, when the cluster size was *t* = 20 and the error probability was *p* = 0.142625, the value of *k*_1_succ_ of Algorithm 5 was 2, while the other algorithms presented *k*_1_succ_ values of at least 12. The results of these simulations for cluster sizes of *t* = 10 and *t* = 20 can be found in Figure 6, Figure 7 and Figure 8.

### 7.2 Results on Data from DNA Storage Experiments

In this section we present the results of the tested algorithms on data from previous DNA storage experiments [20, 24, 40]. The clustering of these data sets was made by the SOLQC tool [45]. We performed each of the tested algorithms on the data and evaluated the edit error rates. Note that in order to reduce the runtime of Algorithm 5 we filtered clusters of size *t >* 25 to have only the first 25 traces. Also here, Algorithm 5 presented the lowest edit error rates in almost all of the tested data sets. These results are depicted in Figure 9.

### 7.3 Performance Evaluation

We evaluated the performance of the different algorithms discussed in this paper. The performance evaluation was performed on our server with Intel(R) Xeon(R) CPU E5-2630 v3 2.40GHz. We implemented our algorithms as well as the previously published algorithm from [23], which presented the second-lowest error rates in our results from Section 7.1. In order to present reliable performance evaluation, the clusters in our experiments were reconstructed in serial order. However, it is important to note, that for practical uses, additional performance improvements can be made by performing the algorithms on the different clusters in parallel and shortening the running time.

- Experiment A included performing reconstruction of simulated data of 2,000 clusters of sequence length of *n* = 100, *p*_*d*_ = *p*_*s*_ = *p*_*i*_ = 0.05, so the total error rate was 0.142625. The cluster sizes were distributed uniformly between *t* = 1 and *t* = 40. The algorithm from [23] (with parameter *w* = 3) reconstructed the full data set in one second with total error rate of 0.03028. Algorithm 5 reconstructed the 2,000 clusters in 2,887 seconds and presented roughly 50% less errors with total edit error rate of 0.014925.
- Experiment B included performing full reconstruction of the data set from [24]. Algorithm 5 reconstructed the full data set in 9, 768 seconds with total edit error rate of 0.00053, while the algorithm from [23] (with parameter *w* = 3) reconstructed the full data set in 50 seconds with total error rate of 0.00081, so it presented approximately 53% more errors compare to Algorithm 5.
- Experiment C included performing reconstruction on 200,000 clusters from the data set of [40]. Algorithm 5 reconstructed the clusters in 456, 200 seconds and presented total edit error rate of 0.00352, while the algorithm from [23] (with parameter *w* = 3) reconstructed the clusters in 296 seconds and total edit error rate of 0.012, which is approximately 3.4 times more errors. Since for this data set, the divider BMA presented the lowest error rate, we decided to evaluate its performance on this data set. The divider BMA algorithm reconstructed the data set in 234 seconds and error rate of 0.0031.

Lastly, to improve the performance of Algorithm 5, we created a hybrid algorithm, that invokes the algorithm from [23] on clusters with more than 20 traces, and otherwise it invokes Algorithm 5. The results of the hybrid algorithm on the simulated data from the experiment A had an edit error rate of 0.02 and in 330 seconds for the first experiment of the simulated data. Since in data from previous DNA storage experiments the variance in the cluster size and in the error rates can be really high [45], we added an additional condition to the hybrid algorithm, so it invokes the algorithm from [23] if the cluster is of size 20 or larger, or if the absolute distance of the difference between the average length of the traces in the cluster and the design length is larger than 10% of the design length. The hybrid algorithm reconstructed the full data set of [24] (experiment B) in 37 seconds and presented error rate of 0.000676. We also performed the hybrid algorithm on 200,000 clusters from the data set of [40] (experiment A). The hybrid algorithm reconstructed these 200,000 clusters in 82, 508 seconds, with edit error rate of 0.002295. The results of the performance experiments are also depicted in Table 1.

**Table 1:**
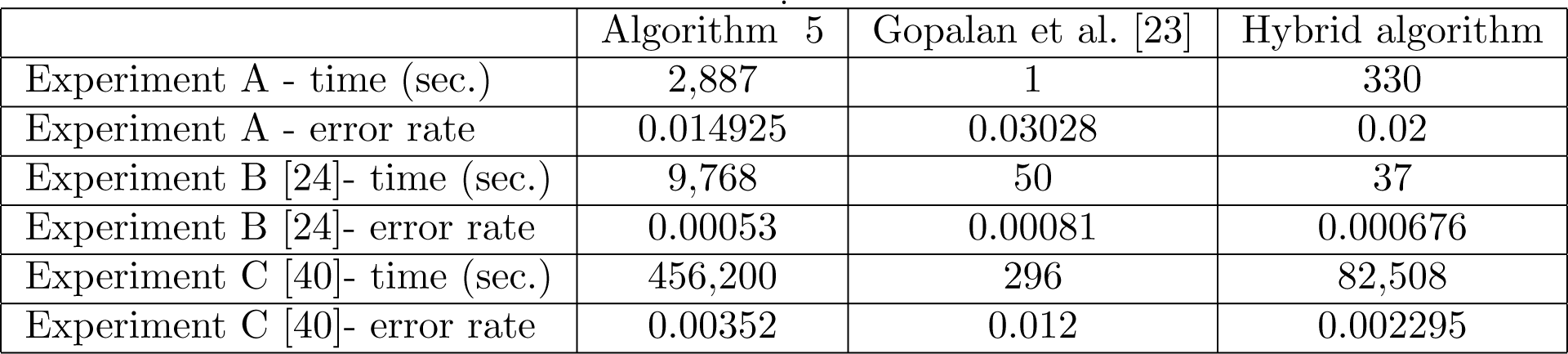
Performance evaluation of Algorithm 5, Gopalan et al. algorithm [23], and the hybrid algorithm. The number presented the running time in seconds and the error rate of each of the algorithms for each of the experiments.

|  | Algorithm 5 | Gopalan et al. [23] | Hybrid algorithm |
| --- | --- | --- | --- |
| Experiment A - time (sec.) | 2,887 | 1 | 330 |
| Experiment A - error rate | 0.014925 | 0.03028 | 0.02 |
| Experiment B [24]- time (sec.) | 9,768 | 50 | 37 |
| Experiment B [24]- error rate | 0.00053 | 0.00081 | 0.000676 |
| Experiment C [40]- time (sec.) | 456,200 | 296 | 82,508 |
| Experiment C [40]- error rate | 0.00352 | 0.012 | 0.002295 |

## 8 Conclusions

We presented in this paper several new algorithms for the deletion DNA reconstruction problem and for the DNA reconstruction problem. While most of the previously published algorithms use a symbol-wise majority approaches, our algorithms look globally on the entire sequence of the traces, and use the LCS or SCS of a given set of traces. Our algorithms are designed to specifically support DNA storage systems and to reduce the edit error rate of the reconstructed sequences. According to our tests on simulated data and on data from DNA storage experiments, we found out that our algorithms significantly reduced the error rates compared to the previously published algorithms. Moreover, our algorithms performed even better when the error probabilities were high, while using less traces than the other algorithms. Even though our algorithms improved previous results, there are still several challenges that need to be addressed in order to fully solve the DNA reconstruction problem. Some of these challenges are listed as follows.

1. Design efficient reconstruction algorithms that improve the current edit error rate.
2. Design error correcting codes for DNA storage systems.
3. Design efficient coded trace reconstruction algorithms for DNA storage systems.
4. Standardization of reconstruction algorithms for DNA storage systems.

## Acknowledgment

The authors thank Matika Lidgi for her help with the divider BMA algorithm and Rotem Samuel for his help with the implementations and simulations of the algorithms in the paper. They also thank Cyrus Rashtchian for helpful discussions. We thank Gala Yadgar for her invaluable help with the simulation infrastructure.

## Footnotes

1 Note that there are many interpretations for the deletion-insertion-substitution, most of them differ on the event when more than one error occured on the same index. Our interpretation of this channel is described in Section 7.1

2 These priorities were selected to support our definition of the deletion-insertion-substitution channel. However, for practical uses, one can easily change these priorities if some preliminary knowledge of the error rates in the data is given.

3 A subsequence can have more than one set of such indices, while the number of such sets is defined as the embedding number [2, 19]. In our algorithm, we chose one of these sets arbitrarily.

4 The parameters we used for the window size *w* of the algorithm from [23] were 2 ⩽ *w* ⩽ 4, and we present the results for all of them.

5 The parameters we used for the algorithm from [52] were ℓ = 5, *δ* = (1 + *p*_*s*_)*/*2, *r* = 2 and *γ* = 3*/*4, while for the data of the DNA storage experiments the substitution probability *p*_*s*_ was taken from [45].

